# Anatomical and Molecular Characterization of the Zebrafish Meninges

**DOI:** 10.1101/2025.04.09.646894

**Authors:** Marina Venero Galanternik, Daniel Castranova, Ryan D. Gober, Tuyet Nguyen, Madeleine Kenton, Gennady Margolin, Aurora Kraus, Abhinav Sur, Louis E. Dye, Van Pham, Adilenne Maese, Melanie Holmgren, Aniket V. Gore, Bakary Samasa, Allison Goldstein, Andrew E. Davis, Avery A. Swearer, James Iben, Tianwei Li, Steven L. Coon, Ryan K. Dale, Jeffrey A. Farrell, Brant M. Weinstein

**Affiliations:** Division of Developmental Biology, Eunice Kennedy Shriver National Institute of Child Health and Human Development, Bethesda, MD; Department of Human Genetics, University of Utah School of Medicine, Salt Lake City, UT; Bioinformatics and Scientific Programming Core, Eunice Kennedy Shriver National Institute of Child Health and Human Development, Bethesda, MD; Microscopy & Imaging Core, Eunice Kennedy Shriver National Institute of Child Health and Human Development, Bethesda, MD; Molecular Genomics Core, Eunice Kennedy Shriver National Institute of Child Health and Human Development, Bethesda, MD

**Keywords:** meninges, scRNA-seq, perivascular cells, FGPs, leptomeninges, pachymeninges

## Abstract

The meninges are a set of connective tissue layers that surround the central nervous system, protecting the brain from mechanical shock, supporting its buoyancy, guarding it from infection and injury, and maintaining brain homeostasis. Despite their critical role, the molecular identity, developmental origins, and functional properties of the cell types populating the meninges remain poorly characterized. This is in large part due to lack of cell type specific markers and difficulty in visualizing and studying these structures through the thick mammalian skull. Here, we show that the zebrafish, a genetically and experimentally accessible vertebrate, possesses an easily imaged mammalian-like meninges. Anatomical and cellular characterization of its composition via histology, electron microscopy, and confocal imaging shows that the adult zebrafish possesses complex multilayered meninges with double-layered dura mater and intricate leptomeningeal layers. Using single cell transcriptomics, we define the molecular identities of meningeal cell populations, including a unique *ependymin* (*epd*)-expressing cell population that constitutes the major cellular component of the leptomeningeal barrier and is essential for brain development and survival. These findings support the use of zebrafish as a useful comparative model for studying the meninges, provide a foundational description for future zebrafish meningeal research, and identify a new Leptomeningeal Barrier Cell that serves as the primary epithelial cell component of the leptomeninges.

## INTRODUCTION

The meninges are a multilayered tissue that envelops and protects the central nervous system (CNS). The cephalic meninges, located between the skull and the brain, act as cushions against external mechanical shock, anchoring the brain inside the skull, supporting buoyancy, and maintaining CNS homeostasis (Decimo et al., 2012; Morris et al., 2022; O’Rahilly and Muller, 1986). Meningeal disease or injury is generally life-threatening, responsible for thousands of deaths every year and can result in severe complications such as chronic seizures, blindness, hearing loss, cognitive impairment and various levels of paralysis (Alves de Lima et al., 2020; Rohde, 2012).

In mammals, the meninges are comprised of three distinct layers. The outermost layer, called the dura mater, is a thick membranous tissue located immediately inside the skull with two adjacent sub-layers, the periosteal dura and the meningeal dura, known collectively as the pachymeninges. The dura mater contains dural sinuses, major venous vessels of the dura mater, as well as recently rediscovered meningeal lymphatic vessels (Absinta et al., 2017; Louveau et al., 2015). The arachnoid and pia mater meningeal layers (known together as the leptomeninges) are found immediately below the dura mater and above the brain parenchyma (Absinta et al., 2017; Dasgupta and Jeong, 2019; Louveau et al., 2015; Morris et al., 2022; Weller et al., 2018). The meninges are populated by numerous types of immune cells, which play roles in the onset of neurodegenerative diseases such as Alzheimer’s and Parkinson’s and have been under increased functional exploration in recent years (Brioschi et al., 2021; Cugurra et al., 2021; Rustenhoven et al., 2021). These studies have focused on the meningeal lymphatics as the most likely anatomical route for immune cells to traffic to and from the brain (Alves de Lima et al., 2020; Da Mesquita et al., 2018; Kanamori and Kipnis, 2020; Louveau et al., 2018; Papadopoulos et al., 2020; Salvador et al., 2024). Additionally, the meninges are highly vascularized. Meningeal damage – due for example to physical trauma – can easily result in the formation of hematomas, causing inflammation and severe brain pressure (Livingston et al., 2017; Russo et al., 2018).

From a developmental standpoint, meningeal formation is important for proper skull and brain development, although only a few studies addressing these interactions have been reported (Choe et al., 2012; Farmer et al., 2021; Mishra et al., 2016; Siegenthaler et al., 2009). Despite their critical role in brain health and hemostasis, the meninges themselves remain a relatively understudied set of tissues compared to the structure they protect, the CNS. The lack of specific meningeal markers has made *in vivo* analysis of this tissue difficult and the molecular identities, developmental origins, and functional properties of cells residing in the meninges remain largely uncharacterized (Derk et al., 2021).

Compared to mammals, even less is known about the meninges of other vertebrates. Classical anatomical studies performed by Giuseppe Sterzi in the late 1800’s suggested that the number of meningeal layers varies among vertebrates (Sterzi, 1902). Sterzi and other contemporary anatomists reported that reptiles and amphibians possess two meningeal layers, while birds possess three (Kappers, 1926; Sterzi, 1902; Wyneken, 2007; Zajicova, 1975). Examination of basal fishes (cyclostomes and plagiostomes) led to the subsequently widely held idea that fishes in general possess only a single meningeal layer, the “meninx primitiva,” although more than one meningeal layer was reported even in primitive fishes in some classical studies (Kappers, 1924, 1926; Sagemehl, 1884; Sterzi, 1902), and more recent studies showed that lampreys (jawless fish) have multiple meningeal layers (Klika and Zajicova, 1975; Nakao, 1979). Descriptions of the meninges of teleost fishes have been even more inconsistent, possibly due in part to varying and often low-resolution methods applied at different developmental time points and to different meningeal structures, e.g., cephalic versus spinal meninges (Kappers, 1924, 1926). Teleost meninges were first described using morphological studies in *Barbus sp*. in the late 19th century (Sagemehl, 1884; Sterzi, 1902), followed by several other teleost species in the 1920s (Kappers, 1924, 1926). Although some studies suggested the presence of only a single layered meninges, other studies supported the presence of more than one “meninx” in teleosts, including both a defined endomeninx consisting of a pia-arachnoid (leptomeninx) and a thin ectomeninx analogous to the dura (Hoffmann and Schwarz, 1996; Kappers, 1924, 1926; Momose et al., 1988; Sagemehl, 1884; Sterzi, 1902; Wang et al., 1995). With conflicting reports from these classical studies, the gross anatomy of the teleost meninges remains unclear.

The zebrafish has recently become one of the most widely used teleost research models. Highly amenable to *in vivo* high-resolution imaging and to genetic and experimental manipulation, zebrafish are also evolutionarily conserved vertebrates with small embryos and larvae and easily accessible adult brains conveniently covered by only a thin translucent skull (Castranova et al., 2021). The experimental and genetic accessibility of the zebrafish make it a desirable model for investigating organ development and function in general, but the thin skull and convenient superficial location of the brain surface makes this a powerful comparative model for high-resolution optical imaging and experimental manipulation of meningeal cell types and tissues. To date, however, little is known about the structure of the zebrafish meninges.

In this study, we clarify the anatomical structure of the zebrafish meninges and demonstrate its utility as a comparative model for the study of meningeal development and disease. Detailed anatomical and cellular characterization of the zebrafish meningeal layers using histology and electron microscopy shows that adult zebrafish possess a mammalian-like multilayered meninges with complex, bi-layered skull-associated *dura mater* and brain-associated leptomeningeal layers. Using single cell RNA sequencing (scRNA-seq), we profile and identify cells present in the dural meninges and leptomeninges. In addition to immune cells, fibroblasts, pigment cells, pericytes, and the previously reported (and assumed meningeal) perivascular Fluorescent Granular Perithelial cells “FGPs” – aka muLECS and BLECs (Bower et al., 2017; van Lessen et al., 2017; Venero Galanternik et al., 2017) – we have identified a novel leptomeningeal cell population expressing high levels of ependymin *(epd)*, a teleost specific cerebrospinal fluid glycoprotein. Orthologous genes such as *ependymin related 1* (EPDR1/MERP1) have been identified from sea urchins to humans but their functions remain unclear (Hoffmann, 1992; Hoffmann and Schwarz, 1996; Konigstorfer et al., 1989; Rinder et al., 1992; Shashoua, 1991; Suarez-Castillo and Garcia-Arraras, 2007). We cloned the *epd* promoter and used it to generate transgenic reporter lines exquisitely specific for *epd*-expressing cells, allowing us to directly visualize and manipulate these cells in living animals. These *epd-*positive cells comprise the major epithelial component of the leptomeninges, where they are closely associated with leptomeningeal blood vessels and FGPs. We have designated them “Leptomeningeal Barrier Cells” or “LBCs.” They are distinct from but share a developmental lineage with meningeal fibroblasts. Using transgene-mediated LBC ablation we show that loss of LBCs in early larvae results in progressive lethality at juvenile stages of development. Acute ablation of these cells in the adult leptomeninges causes severe meningeal hemorrhage and lethality. This work validates the anatomical, cellular, and molecular similarities between the zebrafish and mammalian meninges and demonstrates the utility of the zebrafish as a new research organism to study this tissue.

## RESULTS

### Adult zebrafish possess complex multilayered cephalic meninges

In mammals, the meninges include both bilayered skull-associated dural meninges and multilayered brain-proximal leptomeninges (**Fig. 1A**). We investigated whether comparable meningeal layers are present in the zebrafish, a widely used teleost model organism. Adult zebrafish have a well-developed brain encased in a thin skull, facilitating high resolution *in vivo* optical imaging of the brain surface (**Fig. 1B,C**). In pigment-deficient *casper* zebrafish the highly prominent optic lobes of the zebrafish brain are readily observed through the virtually transparent skull (**Fig. 1C**). When the mammalian skull is removed from the brain, the leptomeninges remain attached to the brain surface while the dura mater (pachymeninges) stays attached to the skull (Louveau et al., 2015), and we find the same is true for the adult zebrafish (**Fig. 1D**).

**Figure 1.**
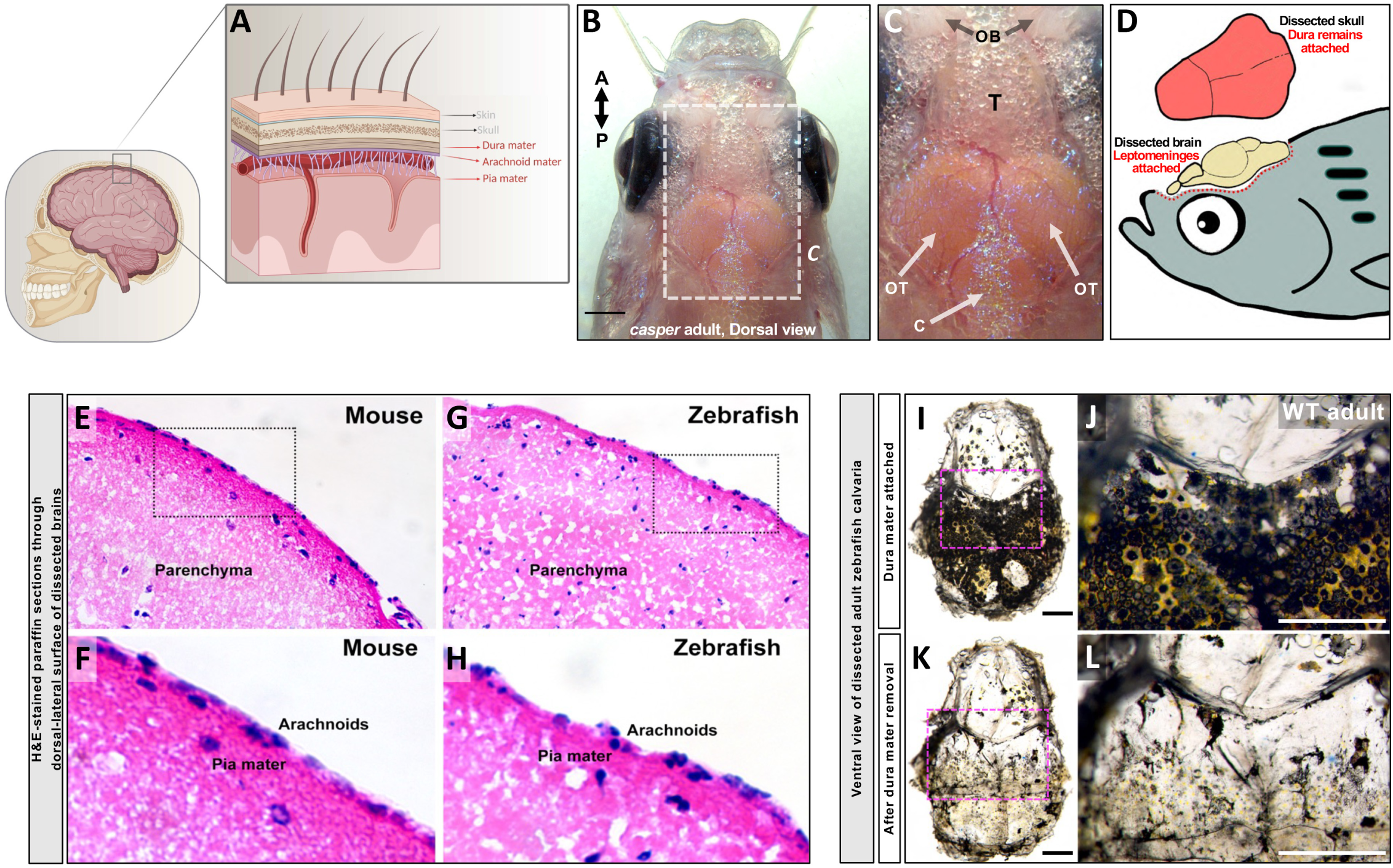
Anatomical characterization of the adult zebrafish cephalic meninges. **A,** Schematic diagram of the structure of the mammalian meninges, including skull-associated dura mater (dural meninges) and deeper arachnoid mater and underlying pia mater immediately proximal to the brain parenchyma (leptomeninges). **B,** Dorsal view of a pigment-deficient adult *casper* zebrafish head (anterior up), with the brain surface visible through the thin transparent skull. **C,** Higher magnification view of the boxed region in panel B, showing olfactory bulbs (OB), telencephalon (T), optic tectum (OT), and cerebellum (C). **D,** Schematic diagram of an adult zebrafish head showing the dissected skull retaining the dura mater and the brain surface retaining the leptomeninges. **E-H,** H&E-stained transverse paraffin sections through the dorsal-lateral surface of dissected mouse (E,F) and zebrafish (G,H) brains showing the brain parenchyma covered by the leptomeninges. Panels F and G show higher magnification images of the boxed areas in E and G, respectively with leptomeningeal pia mater and arachnoids. **I-L,** Dissected calvaria from wild type adult zebrafish, before (I,J) and after (K,L) removal of heavily pigmented dura mater, revealing optically clear skull. Panels K and L show higher magnification images of the boxed areas in I and J, respectively. Scale bars = 1 mm (B), 0.5 mm (I,K), and 1 mm (J,L).

Hematoxylin & Eosin (H&E) sections through the surface of adult mouse (**Fig. 1E,F**) and zebrafish (**Fig. 1G,H**) brains after skull removal reveal similar eosin-dense layers adjacent to the brain parenchyma (**Fig. 1E,G**) surrounded by an outermost haemotoxylin-positive nucleated layer (**Fig. 1F,H**). The ventral side (inside) of the dissected adult zebrafish skull also includes a separate thick membranous layer with large numbers of heavily pigmented cells (**Fig. 1I,J**) that can be removed from the inside of the calvaria, leaving behind a relatively transparent skull (**Fig. 1K,L**). These findings suggest that the zebrafish meninges include multiple cellular layers.

To characterize the anatomy of zebrafish meninges in more detail, we collected transmission electron micrographic (TEM) images of the skull and brain surface, focusing for our initial studies on the optic tectum. Transverse TEM sections through intact zebrafish heads reveal a series of well-defined tissue layers in the space between the zebrafish skull bone and brain including presumptive dura mater, subdural space, and leptomeninges (**Fig. 2A,G**). The subdural space contains extracellular matrix components such as fibrillin and elastin depositions (**Fig. 2A**), which facilitates the manual separation of the leptomeningeal and pachymeningeal layers during dissections. Higher magnification TEM images of sections through the surface of dissected zebrafish brains after skull removal reveal a multilayered epithelial lining separated from the brain parenchyma by a well-defined basal lamina and adjacent glia limitans (**Fig. 2B,C**). These brain-associated leptomeningeal layers (which can be peeled away relatively intact from the parenchyma) are comprised of several cellular sheets as suggested by multiple bilayered membrane-delineated layers (**Fig. 2C**), with numerous blood vessels (**Fig. 2D**) enclosed by non-fenestrated endothelium with abundant and well-defined desmosome-associated tight junctions (**Fig. 2E**), and collagen depositions across the tissue (**Fig. 2F**). TEM of transverse sections through intact zebrafish heads revealed the presence of a defined pachymeninges including distinct and separate periosteal and meningeal dural layers adjacent to the skull (**Fig. 2A,G,H**). Interestingly, we observe characteristic melanophore melanin granules in the meningeal dura (**Fig. 2H**). The presence of melanophores in the dural meninges has also been reported in mammals (Gudjohnsen et al., 2015; Sokolov, 1963). Together, our results suggest that zebrafish possess a complex set of meninges subdivided into multilayered leptomeninges and pachymeninges that resemble the mammalian meninges.

**Figure 2.**
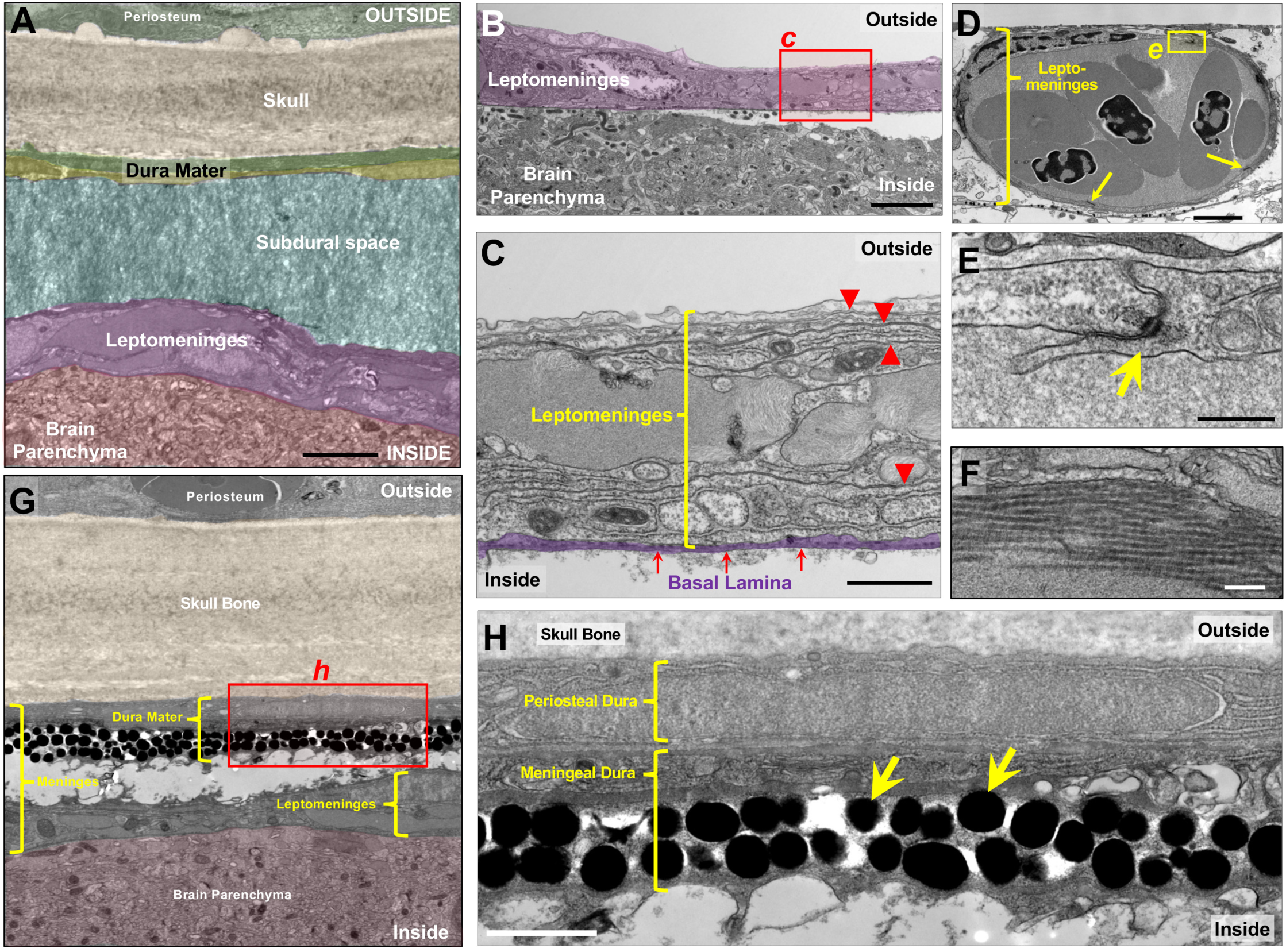
Adult zebrafish possess complex multilayered cephalic meninges. Transmission electron micrograph (TEM) images of transverse sections through the dorsal heads of adult zebrafish, at the level of the optic tecta. **A,** Pseudocolored overview image of the cephalic meninges, revealing multiple tissue layers between the skull and the brain parenchyma. **B,** Dorsal surface of a dissected adult zebrafish brain, showing the leptomeninges partially peeling away from the underlying brain parenchyma. **C,** Higher magnification image of the boxed region in panel B, showing multiple leptomeningeal cell layers separated by membrane bilayers (red arrowheads), delimited ventrally by a well-defined basal lamina (pseudocolored purple). **D,** Nucleated red blood cell-containing blood vessel embedded within the leptomeninges, with desmosomes and junctions connecting individual endothelial cells (arrows). **E,** Higher magnification image of the boxed region in panel D, showing a portion of the vessel wall with a characteristic endothelial cell-cell junction (arrow). **F,** Higher magnification image of collagen fibers in the adult zebrafish leptomeninges. **G,** Pseudocolored overview image of the cephalic meninges, showing separated heavily pigment dura mater (dural meninges) and leptomeninges. multiple tissue layers (pseudocolored) between the skull and the brain parenchyma. **H,** Higher magnification image of the boxed region in panel G, showing bilayered dural meninges with periosteal dura adjacent to the skull with an underlying meningeal dural cell layer with numerous large melanophore pigment granules (arrows). Scale bars = 3 µm (A), 4 µm (B,G), 1 µm (C,H), 2 µm (D), 0.3 µm (E), and 0.2 µm (F).

### Single-cell characterization of the adult zebrafish meninges

The presence of cells with distinct nuclear and cytoplasmic morphologies in TEM images suggests that the leptomeninges and pachymeninges of the zebrafish are likely populated by a variety of different cellular populations. To help identify some of the cell types populating the adult zebrafish meninges, we performed single cell transcriptomics (scRNA-seq) on cell suspensions prepared from 6-month-old adult (i) pachymeninges collected from the ventral face of freshly dissected zebrafish calvaria, and (ii) leptomeninges peeled from the surface of freshly dissected brains (**Fig. 3A**). Approximately 65% of sequenced dural meningeal cells were initially found in 8 closely related clusters enriched for expression of ribosomal and mitochondrial genes (**Supp. Fig. 1**). Since enriched expression of ribosomal and mitochondrial genes in scRNA-seq clusters is a common indicator of dead or dying cells, we bioinformatically filtered these cells out from our pachymeningeal (dural meningeal) single cell data set (**Supp. Fig. 1**) and re-clustered the remaining cells to generate the final pachymeningeal cell clusters shown in **Fig. 3B**. When leptomeninges were manually removed from the brain surface some underlying brain parenchymal tissue invariably adhered to portions of the dissected membranes Indeed, 75% of cells from our leptomeningeal scRNA-seq clustered as readily identifiable neural cell populations (**Supp. Fig. 2A,B**). The neuronal cells could be subdivided into GABAergic and glutamatergic neurons (**Supp. Fig. 2C**). Perhaps unsurprisingly given the dorsal surface location of the habenula in the brain and the fact that it is surrounded by the meninges in mammals (Eckenrode et al., 1992; Han et al., 2017), habenular neurons were very well represented within this group, and could be further separated into medial (MHb) and lateral (LHb) habenular subpopulations (**Supp. Fig. 3**). Since we have not observed cells expressing neuron-specific transgenes in the zebrafish leptomeninges, we bioinformatically filtered most of the neuronal cells from our leptomeningeal single cell data set and the remaining cells were re-clustered to generate the final leptomeningeal cell clusters shown in **Fig. 3C**.

**Figure 3.**
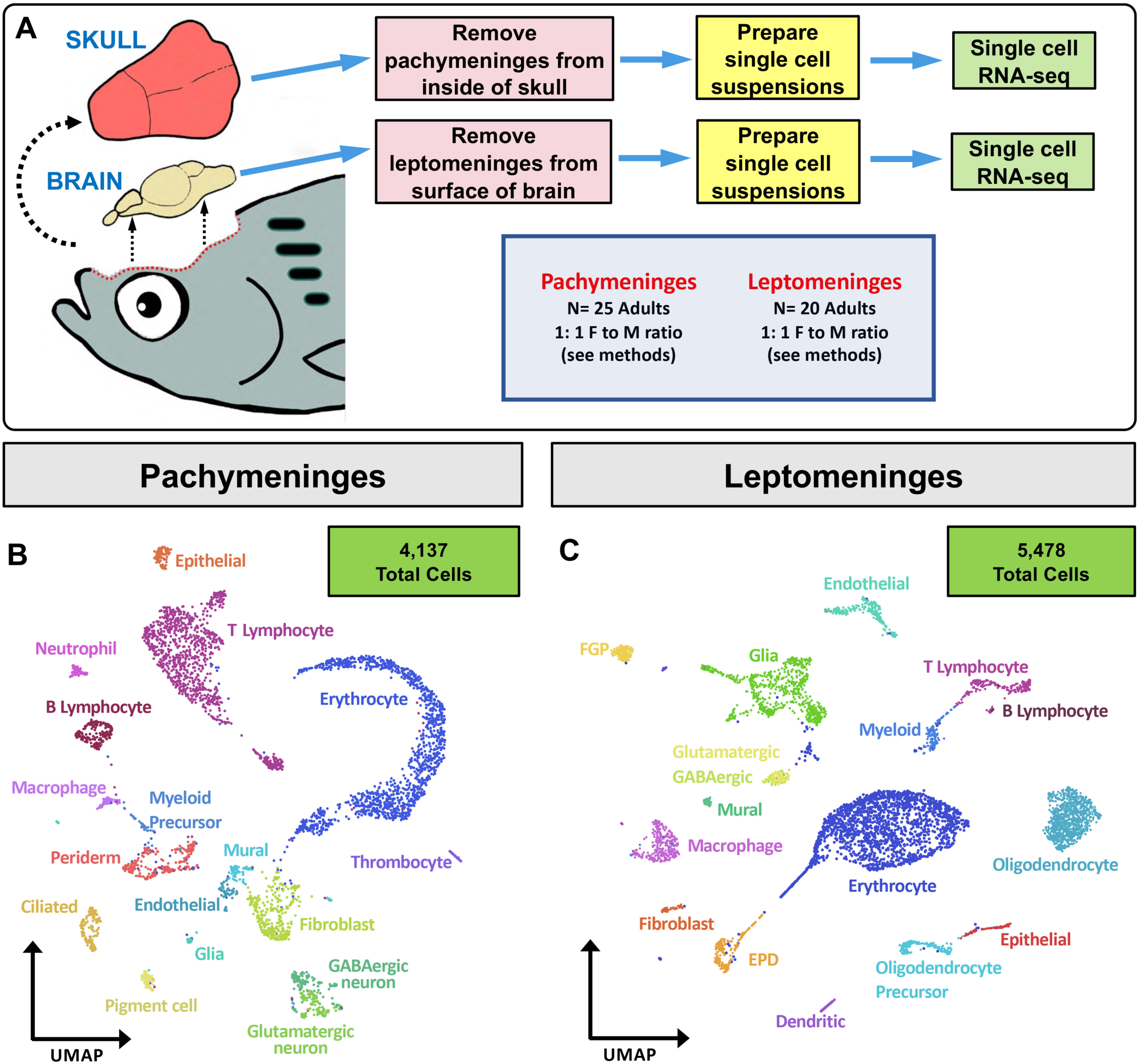
Single-cell characterization of the adult zebrafish meninges. **A,** Schematic diagram of workflow for single cell RNA sequencing (scRNA-seq) of the adult zebrafish meninges. **B,C,** Uniform Manifold Approximation and Projection (UMAP) plots of scRNA-seq data obtained for dissected pachymeninges (dural meninges) (B) and leptomeninges (C) after filtering (see methods and Supp. Figures), showing identities of cell clusters.

### Endothelial and vascular-associated cells in the zebrafish meninges

As noted above, the mammalian meninges are highly vascularized tissues, and the zebrafish meninges similarly contain elaborate vascular networks. Cell clusters enriched for characteristic blood endothelial cell (BEC) markers such as *cdh5, kdrl,* and *cldn5a* (Lawson and Weinstein, 2002; Sauteur et al., 2014; Venero Galanternik et al., 2017) and mural cell markers such as *pdgfrb* and *notch3* (Hellstrom et al., 1999; Stratman et al., 2017; Wang et al., 2014) were detected in the zebrafish leptomeningeal scRNA-seq data set (**Fig. 4A,B**). Confocal imaging of dissected leptomeninges from *Tg(kdrl:mcherry)^y206^*, *Tg(pdgfrb:eGFP)^ncv22^* double transgenic adult zebrafish reveals a dense network of kdrl-positive blood vessels covered by highly ramified, intricate pdgfrb-positive mural cells (**Fig. 4C,D, Supp Fig. 4**). Lymphatic vessels are not present in the mammalian leptomeninges, and we have not observed them in the leptomeninges of adult zebrafish, but a cell cluster is present in the leptomeningeal scRNA-seq data set that expresses characteristic lymphatic endothelial cell (LEC) markers such as *mrc1a, lyve1b, stab1, stab2,* and *flt4* (**Fig. 4B**). These cells are an unusual perivascular cell population that we and others previously reported on the outer surface of the zebrafish brain (Bower et al., 2017; van Lessen et al., 2017; Venero Galanternik et al., 2017) that strongly resemble mammalian FGPs, perivascular macrophage-like cells described in the leptomeninges of mice, rats, and humans (Mato et al., 1986; Mato and Mato, 1983; Mato et al., 1981; Mato et al., 1982; Ookawara et al., 1996). Despite appearing as individual separated cells rather than part of a tube, and despite their macrophage-like cell morphology and perivascular location, zebrafish FGPs (also referred to as Mural Lymphatic Endothelial Cells, “muLECs” or Brain Lymphatic Endothelial Cells, “BLECs”) have a molecular signature very similar to lymphatic endothelial cells and they arise from primitive lympho-venous endothelium (van Lessen et al., 2017; Venero Galanternik et al., 2017). High-resolution confocal imaging of the meninges of *Tg(kdrl:mcherry)^y206^*, *Tg(mrc1a:egfp)^y251^* transgenic animals shows that, as in mammals, zebrafish FGPs are exclusively localized to the inner, brain-associated leptomeninges in close association with leptomeningeal blood vessels (**Fig. 4E,F**). FGPs have well-documented scavenger properties with large autofluorescent vesicles (ie, “fluorescent granules”) that accumulate substances taken up from the extracellular environment including infiltrated dyes such as Dextran or India ink (Huisman et al., 2022; van Lessen et al., 2017; Venero Galanternik et al., 2017). Although as noted above lymphatic vessels are not present in the leptomeninges or the brain proper in mammals, they are found in the more superficial skull-associated dural meninges (Aspelund et al., 2015; Louveau et al., 2015). High-resolution confocal imaging through the skull of intact, living *Tg(kdrl:mcherry)^y206^*, *Tg(mrc1a:egfp)^y251^*double-transgenic zebrafish adults injected intravascularly with Draq5 to mark circulating blood cells shows that the skull-associated dural meninges (pachymeninges) contains both kdrl-positive blood vessels with robust blood flow and *bona fide* mrc1a-positive lymphatic vascular tubes lacking erythrocytes (**Fig. 4G,H**, (Castranova et al., 2021)). The presence of these structures in the pachymeninges is also reflected in our scRNA-seq data, which includes cell clusters expressing characteristic BEC, mural cell, and LEC markers (**Fig. 4I**).

**Figure 4.**
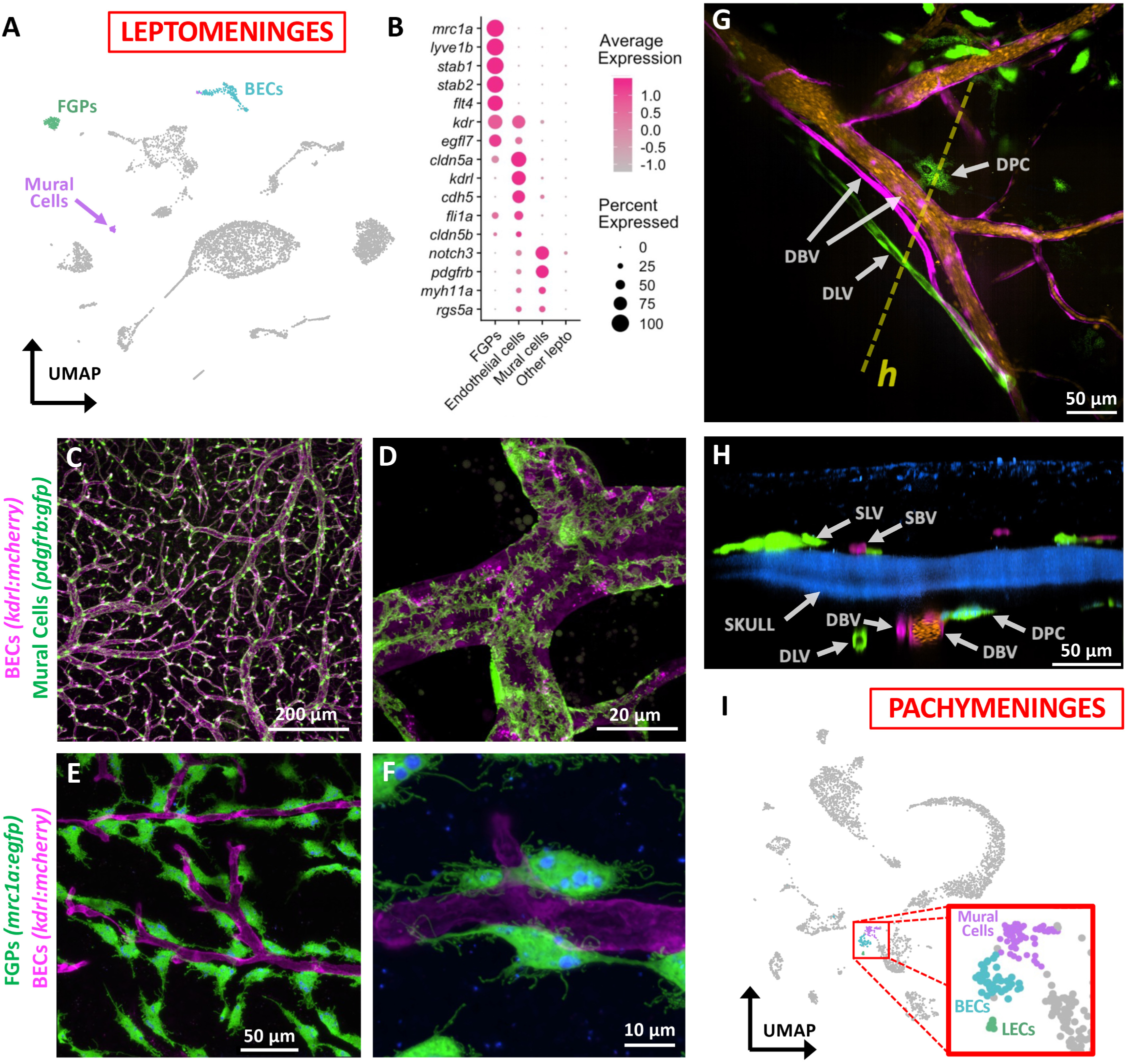
Endothelial and vascular-associated cells in the zebrafish meninges. **A,** UMAP plot of leptomeningeal scRNA-seq data with blood endothelial cell (BEC), mural cell, and Fluorescent granular perithelial cell (FGP) cell clusters highlighted. **B,** Dot plot showing enriched expression of selected diagnostic genes in FGPs, BECs, and mural cells. Average scaled expression and percent of cells in each cluster expressing each gene is indicated by dot color and size, respectively. **C,D,** Confocal images of leptomeningeal blood vessels in *Tg(kdrl:mcherry)^y206^*, *Tg(pdgfrb:egfp)^ncv22^* double transgenic adult zebrafish with kdrl:mcherry-positive BECs (magenta) and pdgfrb:GFP-positive mural cells (green). Panel C shows lower magnification overview, panel D shows higher magnification image revealing complex, branched mural cells covering vessels. **E,F,** Confocal images of leptomeningeal blood vessels in *Tg(kdrl:mcherry)^y206^*, *Tg(mrc1a:egfp)^y251^* double transgenic adult zebrafish with kdrl:mcherry-positive BECs (magenta) and mrc1a:egfp-positive FGPs (green) containing characteristic autofluorescent internal vesicles (blue). Panel E shows lower magnification overview, panel F shows higher magnification image of distinct, separated FGPs closely apposed to blood vessels. **G,H,** Confocal images of the dural meninges in *Tg(kdrl:mcherry)^y206^*, *Tg(mrc1a:egfp)^y251^* double transgenic adult zebrafish with intravascular Draq5 labeling, showing kdrl:mcherry-positive blood vessels (magenta), Draq5 labeled blood vessel lumens (orange), mrc1a:egfp-positive lymphatic vessels (green) and autofluorescent dural pigment cells (green), and skull autofluorescence (blue). Panel G is a dorsal view and panel H is a lateral view of the same confocal stack. The dashed yellow line in panel G shows the plane of section viewed in panel H. **I,** UMAP plot of dural meningeal (pachymeningeal) scRNA-seq data with mural cell, blood endothelial cell (BEC), and lymphatic endothelial cell (LEC) cell clusters highlighted. Abbreviations: DBV, dural blood vessel, DLV, dural lymphatic vessel, DPC, dural pigmented cell, SBV, skin blood vessel, SLV, skin lymphatic vessel. Scale bars = 200 µm (C), 20 µm (D), 50 µm (E), 10 µm (F), 50 µm (G), and 50 µm (H).

### Hematopoietic cells in the zebrafish meninges

The mammalian meninges play a key role in nourishing and protecting the brain, and they contain abundant numbers of blood cells that participate in carrying out these functions. Zebrafish leptomeningeal and pachymeningeal (dural) scRNA-seq reveals similar large numbers of hematopoietic cells (**Fig. 3B,C**). Erythrocytes represent the largest cell population identified in both data sets – 1,861 (33.9%) and 1,420 (34.3%) in leptomeninges and dura respectively, as shown by highly enriched expression of a variety of definitive erythrocyte-specific transcripts, including *hbba1, hbba1.1, hbaa1, hbba2* and *hbaa2* hemoglobin genes (**Supp. Fig. 5**). Erythrocytes with characteristic morphology could also be readily observed within blood vessels in TEM and confocal images of the meninges (e.g., **Fig. 2D** - like camels, fish have nucleated erythrocytes).

The mammalian meninges also contain an array of different immune cell populations that mediate active immune surveillance to help protect the brain from infection (Brioschi et al., 2021; Cugurra et al., 2021; Rustenhoven et al., 2021). In the zebrafish, the leptomeninges and dural meninges contain a similar variety of immune cell types, including T cells, B cells, and macrophages (**Fig. 5A-D**). The presence of these immune cell populations is confirmed by high-resolution confocal imaging of the meninges in zebrafish carrying transgenes that mark specific immune cell populations, including *Tg(lck:egfp)^cz1^* T cells (**Fig. 5E-G**), *Tg(cd79b:egfp)^fcc89^*B cells (**Fig. 5H**), *Tg(mpeg1:egfp)^gl22^*macrophages (**Fig. 5I-K**), and, rarely, *Tg(lyz:dsred2)^nz50^* neutrophils (**Fig. 5L**). Confocal imaging through intact skulls of living *Tg(lck:mcherry)^ns107^, Tg(mrc1a:eGFP)^y251^* double transgenic animals shows that patrolling T cells are present in the meninges (**Fig. 5M-O, Supp. Movie 1**) Together, immune cell clusters make up 720/5,478 (13.1%) of leptomeningeal and 1,414/4,137 (34.2%) of dural cells captured in our scRNA-seq samples, a major fraction of the total cells present. This suggests that, as in mammals, the zebrafish meninges play an important role in immune surveillance and protection of the brain.

**Figure 5.**
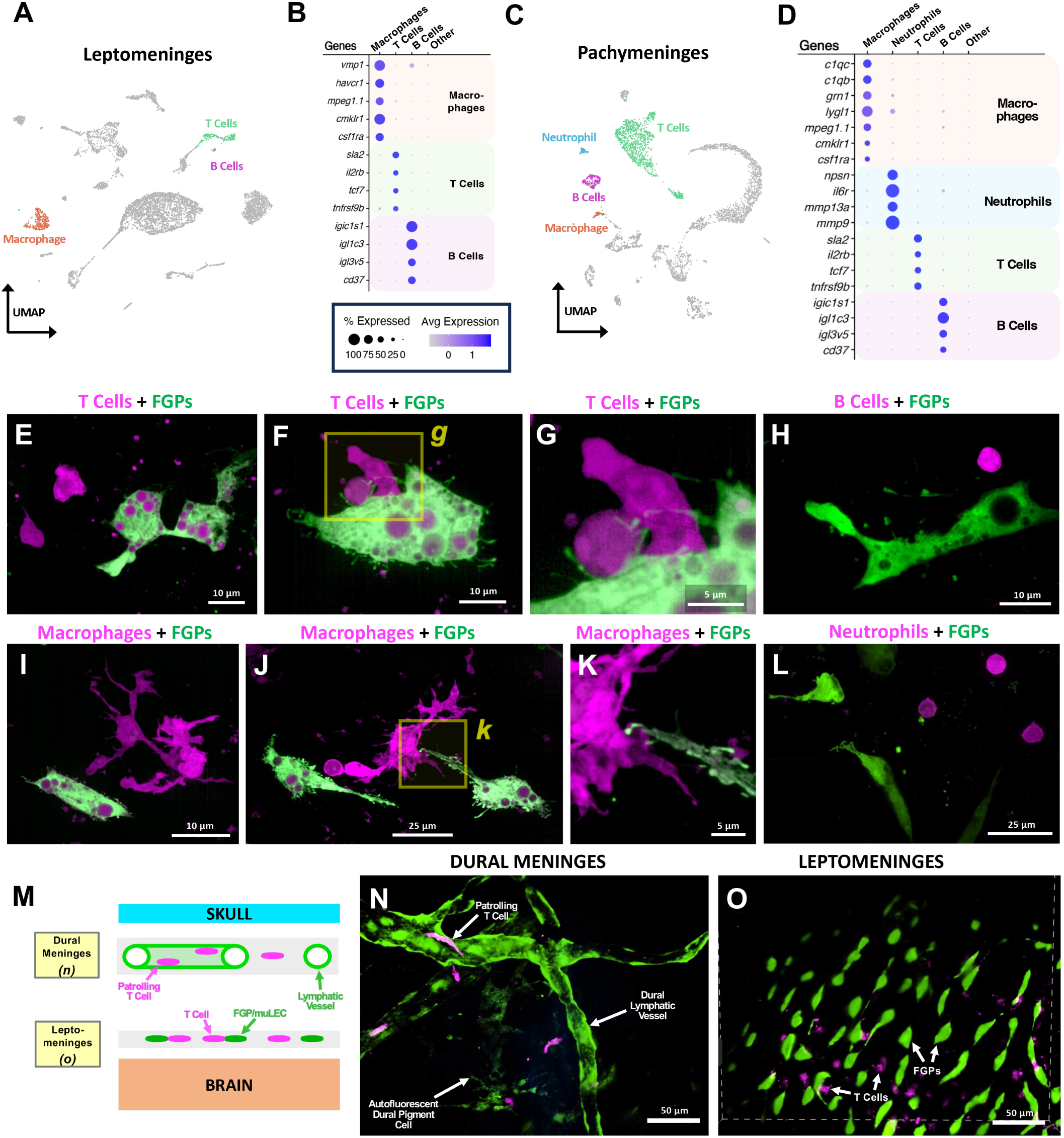
Hematopoietic cells in the zebrafish meninges. **A,** UMAP plot of leptomeningeal scRNA-seq data with T cell, B cell, and macrophage cell clusters highlighted. **B,** Dot plot showing enriched expression of selected diagnostic genes in T cells, B cells, and macrophages. **C,** UMAP plot of leptomeningeal scRNA-seq data with T cell, B cell, macrophage, and neutrophil cell clusters highlighted. **D,** Dot plot showing enriched expression of selected diagnostic genes in T cells, B cells, macrophages, and neutrophils. For dot plots in panels B and D average scaled expression and percent of cells in each cluster expressing each gene is indicated by dot color and size, respectively. **E-L,** Representative confocal images of cells in the adult leptomeninges of animals expressing different immune cell type-specific transgenes (magenta), including: (**E-G**) *Tg(lck:egfp)^cz1^* T cells (magenta), (**H**) *Tg(cd79b:egfp)^fcc89^*B cells (magenta), (**I-K**) *Tg(mpeg1:egfp)^gl22^* macrophages (magenta), and, rarely, (**L**) *Tg(lyz:dsred2)^nz50^* neutrophils (magenta). Leptomeningeal FGPs (green) are also observed with the animals in panels E-K are also co-expressing the *Tg(lyve1b:dsred)^nz101^* transgene and the animals in panel L also co-expressing the *Tg(mrc1a:eGFP)^y251^* transgene (both shown in green, with magenta autofluorescent internal vacuoles). The images in panels G and K are higher magnifications of the boxed regions in panels F and J, respectively. **M,** Schematic diagram noting the location of patrolling T cells and lymphatic vessels in the dural meninges, and T cells and FGPs in the leptomeninges. **N,O,** Selected planes from different levels of a confocal image stack taken through the dorsal head of *Tg(lck:egfp)^cz1^*, *Tg(lyve1b:dsred)^nz101^*double transgenic animals, showing T cells (magenta) in and around lymphatic vessels (green) in the dural meninges (panel N) or adjacent to FGPs in the leptomeninges (panel O). Scale bars = 10 µm (E,F,H,I), 5 µm (G,K), 25 µm (J,L), and 50 µm (N,O).

### A novel *ependymin*-expressing Leptomeningeal Barrier Cell

The zebrafish leptomeningeal scRNA-seq data set also contains a unique cluster of cells expressing very high levels of *ependymin* (*epd*), a teleost cerebrospinal fluid glycoprotein previously implicated in memory, neural regeneration and dominance in studies performed in carp, goldfish and salmonids (Hoffmann, 1994; Piront and Schmidt, 1988; Schmidt et al., 1991; Schmidt et al., 1995; Shashoua, 1988, 1990, 1991; Suarez-Castillo et al., 2004; Tang et al., 1999). Although these cells share some markers with meningeal fibroblasts (DeSisto et al., 2020), such as *FXYD domain containing ion transport regulator 1 (phospholemman; fxyd1), fatty acid binding protein 11a (fabp11a),* and *insulin-like growth factor binding protein 2a (igfbp2a),* they cluster as a distinct and separate cell population in our leptomeninges scRNA-seq data set (**Fig. 6A,B**; see below).

**Figure 6.**
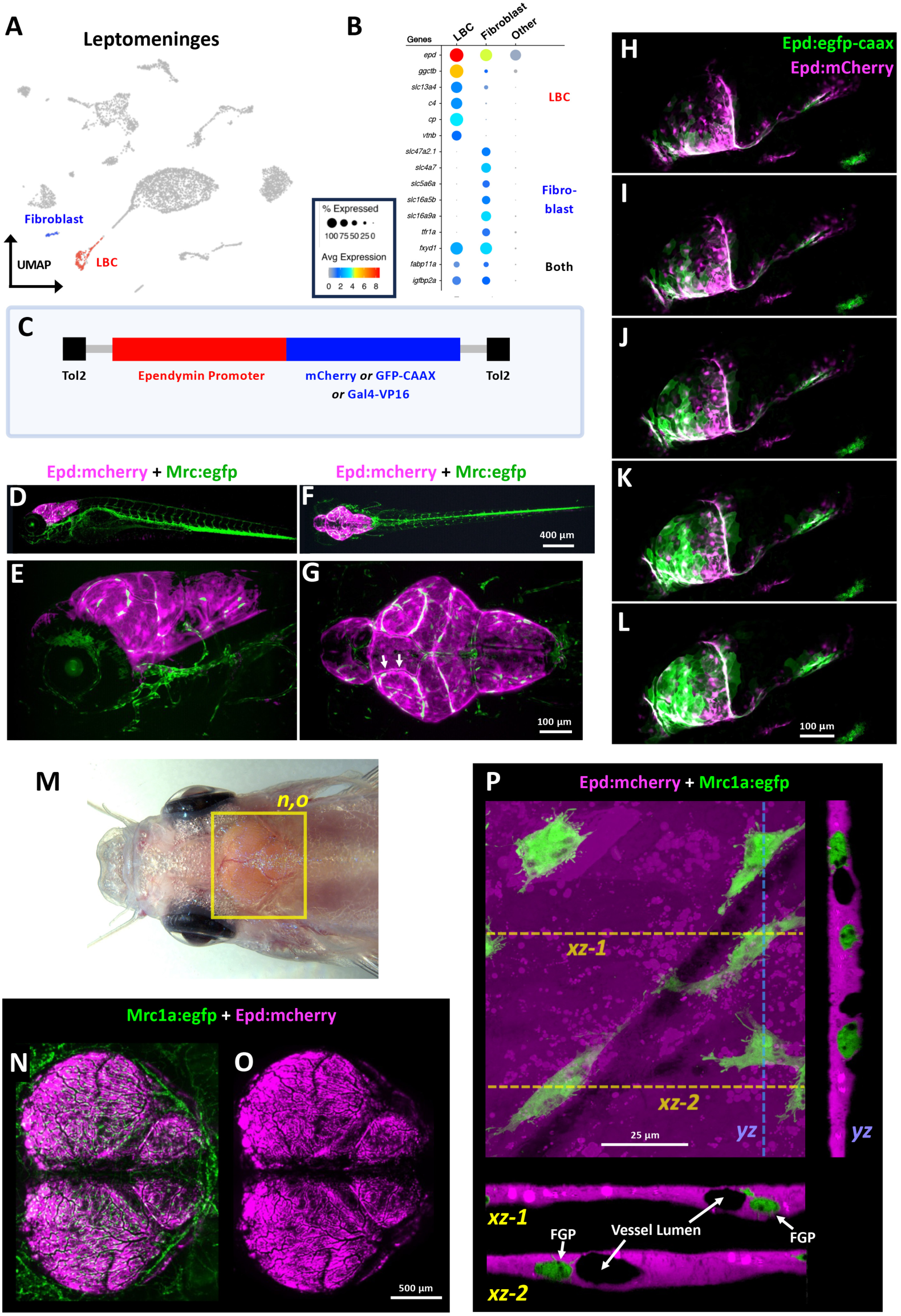
LBCs, a novel *ependymin*-expressing leptomeningeal cell population. **A,** UMAP plot of leptomeningeal scRNA-seq data with meningeal fibroblast (blue) and LBC (red) cell clusters highlighted. **B,** Dot plot showing enriched expression of selected diagnostic genes in meningeal fibroblasts, LBCs, and all other leptomeningeal cells. Average expression and percent of cells in each cluster expressing each gene is indicated by dot color and size, respectively, with average expression on a log2 scale. **C,** Schematic diagram of Tol2 constructs used to generate stable germline *epd* promoter-dependent transgenic zebrafish expressing mCherry, GFP-CAAX, or Gal4-VP16. **D-G,** Lateral (D,E) or dorsal (F,G) view confocal images of a 5 dpf *Tg(epd:mcherry)^y716^*, *Tg(mrc1a:egfp)^y251^*double transgenic animal, showing epd:mcherry expression (magenta) restricted exclusively to the cephalic meninges. Panels E and G show higher magnification images of the head of the same animal shown in panels D and F, respectively. Expression of mrc1a:eGFP (green) marks cephalic FGPs veins, and lymphatics. **H-L,** Confocal image time series of LBC emergence in the developing cephalic meninges of *Tg(epd:mcherry)^y715^*, *Tg(epd:gfp-caax)^y716^*double transgenic zebrafish beginning at 54 hpf (see **Movie 2** for complete series). **M,** Dorsal view of an adult *casper* zebrafish head with the optic tectum and cerebellum clearly visible through the thin, translucent skull. The yellow box shows the approximate region depicted in panels N and O. **N,O,** Confocal images of the meninges imaged through the intact skull of a living *Tg(mrc1a:egfp)^y251^, Tg(epd:mcherry)^y715^* double transgenic adult zebrafish, showing (N) mrc1a:egfp-positive FGPs and lymphatics (green) and epd:mcherry-positive LBCs (magenta), or (O) only epd:mcherry-positive LBCs (magenta). **P,** High-resolution, high magnification confocal Z-stack through the leptomeninges on the surface of a dissected *Tg(mrc1a:egfp)^y251^, Tg(epd:mcherry)^y715^* double transgenic adult zebrafish brain, showing LBCs (magenta) and FGPs (green). The large center image is a dorsal XY view of the reconstructed image stack, and the images below and to the right show single-plane XZ and YZ views along the planes noted by the dashed lines (xz-1, xz-2, yz). See also accompanying **Movie 3**. Scale bars = 400 µm (D,F), 100 µm (E,G), 500 µm (N,O), 100 µm (H-L), and 25 µm (P).

To learn more about these cells, we generated reporter lines using 5 Kb of genomic DNA upstream from the start site of the *ependymin* gene (**Fig. 6C**) to drive expression of transgenes assembled using Tol2kit/Gateway technology (Kwan et al., 2007). Germline transgenic lines generated with *epd* promoter vectors exhibit strong, highly specific expression in the cephalic leptomeninges of both larval (**Fig. 6D-L**) and adult (**Fig. 6N-Q**) zebrafish. In 5 days post fertilization (dpf) *Tg(epd:mcherry)^y716^*, *Tg(mrc1a:egfp)^y251^*double transgenic animals mCherry is expressed exclusively in large, flat cells lining the outside surface of the brain (**Fig. 6D-G, Supp. Movie 2**) whose cell bodies showing significant alignment with major meningeal blood vessels (**Fig. 6G**, arrows). Cranial expression of *Tg(epd:mcherry)^y716^* or *Tg(epd:gfp-caax)^y716^* transgenes is first noted at around 54-60 hours post fertilization (hpf) in cells in the ventral-lateral head that then migrate dorsally over and around the surface of the brain (**Fig. 6H-L, Supp. Movie 2**). In adult zebrafish ependymin-expressing cells also localize exclusively to the leptomeninges. Confocal imaging of the surface of dissected *Tg(mrc1a:egfp)^y251^, Tg(epd:mcherry)^y715^* double-transgenic adult zebrafish brains shows that these cells completely cover the surface of the optic tecta and cerebellum, in close association with FGPs and leptomeningeal blood vessels (**Fig. 6M-O**). Higher magnification confocal imaging of the surface of dissected *Tg(mrc1a:egfp)^y251^, Tg(epd:mcherry)^y715^* double-transgenic adult zebrafish brains shows that they represent the major population of resident epithelial cells in the brain-associated leptomeninges, with leptomeningeal blood vessels and FGPs embedded within and surrounded by layers of these cells (**Fig. 6P, Supp. Movie 3**). Based on their localization and morphology, we refer to these strongly *ependymin*-positive leptomeningeal cells as “Leptomeningeal Barrier Cells” or “LBCs” throughout the rest of this study.

### LBCs are closely related to but distinct from meningeal fibroblasts

To further explore the identity and cellular origins of LBCs and their relationship to meningeal fibroblasts, we mapped the cellular trajectory of the early LBC developmental lineage using “Daniocell” (Farrell et al., 2018; Sur et al., 2023), a comprehensive whole-animal scRNA-seq data set comprised of closely spaced time points encompassing the first 5 days of zebrafish development. Later Daniocell time points contain a mesenchymal cell cluster whose gene expression profile strongly resembles the adult zebrafish leptomeninges LBC cell cluster (“mese.29: meninges-leptomeninges”), as well as a closely related but separate mesenchymal cluster corresponding to meningeal fibroblasts (“mese.32: meninges-meningeal fibroblasts”), with both clusters appearing to differentiate from an earlier-appearing meningeal precursor (“mese.21: meninges-precursors”) cell cluster (**Fig. 7A,B, Supp. Fig. 6**). The Daniocell and adult leptomeninges LBC clusters both have comparable highly enriched expression of diagnostic genes such as *ependymin (epd), gamma-glutamylcyclotransferase b (ggctb), solute carrier family 13 member 4 (slc13a4), complement component 4 (c4), ceruloplasmin (cp),* and *vitronectin b (vtnb)* (**Figs. 6B, 7CD, Supp. Fig. 7A**). The meningeal fibroblast cell clusters in Daniocell and adult leptomeninges scRNA-seq datasets also share enriched expression of a number of genes, notably including several *solute carrier family (slc)* genes including *slc47a2.1, slc4a7, slc5a6a, slc16a5b,* and *slc16a9a,* as well as the *transferrin receptor 1a (tfr1a)* (**Figs. 6B, 7C,D, Supp. Fig. 7A**), many of which are also present in mammalian meningeal fibroblasts (DeSisto et al., 2020). In addition to the genes noted above that are highly expressed in only one of the two cell types, there are also a number of genes that are highly expressed in both cell types in Daniocell and adult leptomeninges scRNA-seq datasets, including the *fxyd1, fabp11a,* and *igfbp2a* genes noted above (**Figs. 6B, 7C,D, Supp. Fig. 7A**). Pseudotime analysis of the Daniocell data set identified a branchpoint from meningeal mesenchymal precursors to LBCs and meningeal fibroblasts around 48-58 hpf (**Supp. Fig. 7B**), approximately the same time that we begin to detect mCherry expression in *Tg(epd:mcherry)^y715^* transgenic animals (**Fig. 6H, Supp. Movie 2**). We then used our trajectories to identify transcription factors that were differentially expressed between the LBCs and meningeal fibroblasts (**Fig. 7E**), changed during their development, and were not broadly expressed across most cell types in the animal. We identified *foxl2a*, which specifically is activated in precursors and peaks as they adopt either LBC or meningeal fibroblast transcriptional states, *zeb2b* which is activated in LBCs as they become transcriptionally distinct, *klf2a* and *six1a* which are expressed in precursors but remain expressed only in meningeal fibroblasts, and *klf15* and *bhlhe40* which seem to be primarily activated in meningeal fibroblasts as they become transcriptionally distinct (**Fig. 7F**). Future functional assays will identify whether these TFs control key aspects of the gene regulatory networks of these cell types.

**Figure 7.**
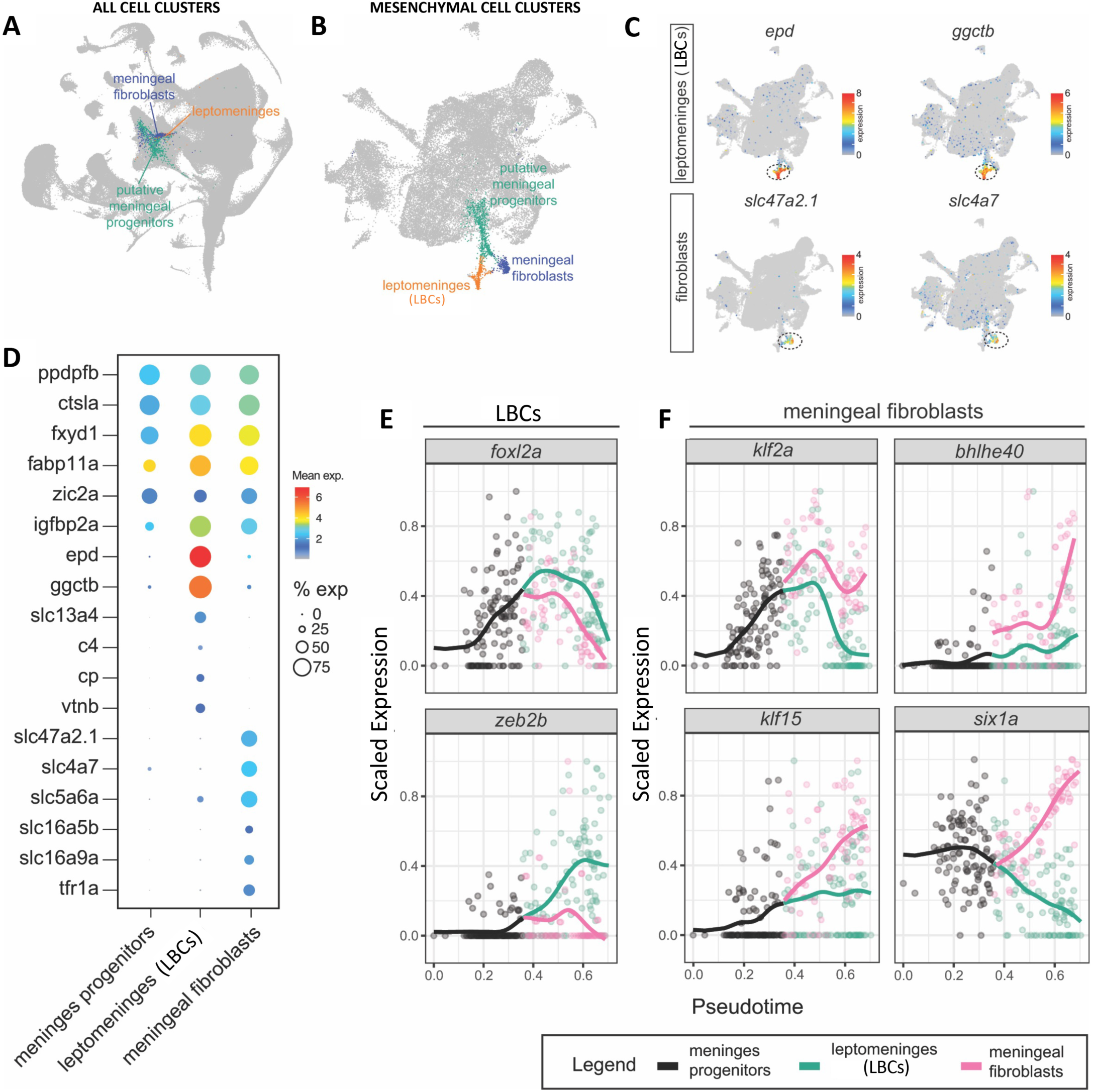
LBCs are closely related to but distinct from meningeal fibroblasts. UMAP plot of all cell clusters in the Daniocell (https://daniocell.nichd.nih.gov/) scRNA-seq dataset for developing zebrafish embryos from the 1-cell stage to 5 dpf, highlighting cell clusters corresponding to meningeal fibroblasts (blue), LBCs (orange) and putative meningeal progenitors (green). **B,** UMAP plot of all mesenchymal cells present in the Daniocell scRNA-seq dataset, with meningeal fibroblast (blue), LBCs (orange) and putative meningeal progenitors (green) clusters highlighted as in (A). **C,** Feature plots of genes differentially expressed in LBCs (*epd, ggctb*) or meningeal fibroblasts (*slc47a2.1, slc4a7*) in the mesenchymal cells represented in panel B. **D,** Dot plot showing enriched expression of selected diagnostic genes in meningeal fibroblasts, LBCs, or both in the mesenchymal subset of the Daniocell scRNA-seq data set. Average expression and percent of cells in each cluster expressing each gene are indicated by dot color and size, respectively, with average expression on a log2 scale. **E,F,** Pseudotime trajectory analysis of transcription factors initially expressed in meningeal mesenchymal progenitors whose expression becomes preferentially enriched in either LBCs (E) or meningeal fibroblasts (F).

### Essential early function of the zebrafish meninges demonstrated by LBC ablation

To examine the functional role of LBCs and the zebrafish meninges as a whole, we generated *Tg(epd:gal4-VP16)^y717^* transgenic zebrafish (**Fig. 6C**) and crossed them to *Tg(uas:ntr-mcherry)^c264^*, *Tg(mrc1a:egfp)^y251^* double transgenic animals to generate *Tg(epd:gal4-VP16)^y717^*, *Tg(uas:ntr-mcherry)^c264^*, *Tg(mrc1a:egfp)^y251^* animals heterozygous for all three transgenes. These fish were intercrossed to generate *Tg(epd:gal4-VP16)^y717^*, *Tg(uas:ntr-mcherry)^c264^*, *Tg(mrc1a:egfp)^y251^* triple homozygous and *Tg(epd:gal4-VP16)^y717^*, *Tg(mrc1a:egfp)^y251^* double homozygous (*uas:ntr-mcherry* negative control) animals for Metronidazole-(MTZ) induced ablation of LBCs. We refer to these below as “NTR” and “GFP only” animals, respectively (**Fig. 8A**). We tested the effects of meningeal LBC ablations in two ways. First, we treated zebrafish larvae with 10 mM MTZ continuously from 5 dpf to 7 dpf to remove differentiated LBCs from the surface of the brain, in order to test whether and for how long developing animals can survive without LBCs or a properly developing leptomeninges (**Fig. 8A-L**). Second, we treated adult zebrafish with 10 mM MTZ for one day to determine the consequences of acute ablation of fully developed LBC-containing leptomeninges in a mature animal (**Fig. 8M-V**).

**Figure 8.**
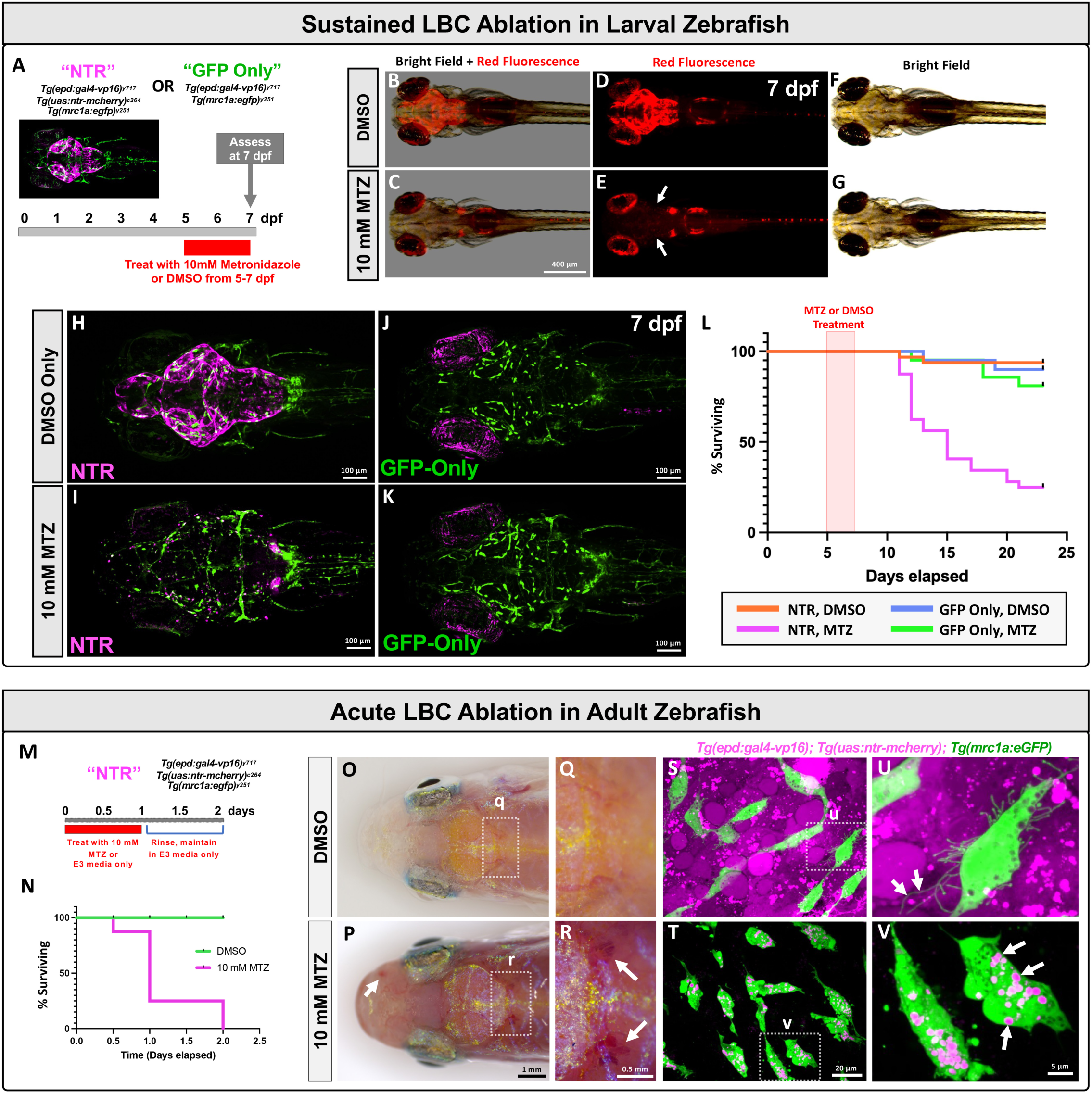
Essential function of the zebrafish meninges demonstrated by LBC ablation. **A,** Schematic diagram of experimental design for LBC ablation in zebrafish larvae, with *Tg(epd:gal4-VP16)^y717^*, *Tg(uas:ntr-mcherry)^c264^*, *Tg(mrc1a:egfp)^y251^* triple transgenic (“NTR”) or *Tg(epd:gal4-VP16)^y717^*, *Tg(mrc1a:egfp)^y251^* double transgenic control (“GFP Only”) animals. Image shows LBCs present at the beginning of treatment. Red bar shows timing of treatment with either 1% DMSO or 10 mM Metronidazole with 1% DMSO. **B-G,** Representative dorsal view bright field and red fluorescence (B,C), red fluorescence only (D,E), and brightfield only (F,G) images of 7 dpf DMSO carrier only (B,D,F) or 10 mM MTZ (C,E,G) treated “NTR” zebrafish larvae. The MTZ-treated animal shows complete ablation of LBC cells (arrows in panel E). **H-K,** Representative dorsal view confocal images of the heads of 7 dpf “NTR” (H,I) or “GFP Only” (J,K) zebrafish larvae treated from 6-7 days with either DMSO carrier only (H,J) or 10 mM MTZ (I,K). The mCherry-positive LBCs are in magenta, and EGFP-positive FGPs and lympho-venous vessels are in green. The NTR-expressing, MTZ-treated animal in panel I shows loss of LBCs as well as strong concomitant reduction in numbers of FGPs compared to controls. **L**, Quantification of survival for “NTR” or “GFP Only” animals treated from 5-7 dpf with either DMSO carrier only or 10 mM MTZ, then monitored for survival until 25 dpf. LBC-ablated (NTR, MTZ) animals show a steep decline in survival beginning at approximately 11 dpf. **M,** Schematic diagram of experimental design for LBC ablation in 6 month old *Tg(epd:gal4-VP16)^y717^*, *Tg(uas:ntr-mcherry)^c264^*, *Tg(mrc1a:egfp)^y251^* triple transgenic (“NTR”) adult zebrafish. Red bar shows timing of treatment with either 10 mM Metronidazole (10 mM MTZ) or DMSO media only, with n=8 fish per condition, 1:1 female to male ratios. **N,** Quantification of survival for adult NTR animals treated with either DMSO carrier only or 10 mM Metronidazole (MTZ). **O-R,** Dorsal images of living NTR adult zebrafish one day after beginning treatment with DMSO only (O,Q) or 10 mM MTZ (P,R). Panels Q and R show higher magnification views of the boxed regions in panels O and P, respectively. MTZ-treated animals showed clear evidence of hemorrhage (white arrows, P,R). **S-V,** Confocal micrographs of the surface of brains freshly dissected from NTR adult zebrafish one day after beginning treatment with either DMSO only (S,U) or 10 mM MTZ (T,V). Panels U and V show higher magnification views of the boxed regions in panels S and T, respectively. The mCherry-positive LBCs are absent from MTZ-treated animals. FGPs in MTZ-treated animals are still present but their morphology is abnormal, and they contain large numbers of strongly mCherry-positive vacuoles (arrows in panel V). Scale bars = 400 µm (B-G), 100 µm (H-K), 1 mm (O,P), 0.5 mm (Q,R), 20 µm (S,T), and 5 µm (U,V).

NTR larvae incubated continuously in 10 mM MTZ from 5 to 7 dpf initially survived LBC ablation at the same rate as GFP-Only animals treated with MTZ or DMSO (**Fig. 8B-G**). However, unlike NTR larvae incubated without drug (**Fig. 8B,D,F**), MTZ treated NTR animals displayed complete loss of LBCs (**Fig. 8C,E, Supp. Movie 4**). The MTZ treated NTR larvae lacking LBCs also showed loss and mislocalization of leptomeningeal FGPs (**Fig. 8I**), while NTR larvae incubated only in DMSO media without drug or GFP-Only animals with or without MTZ treatment retained their FGPs (**Fig. 8H,J,K**). Despite the general healthy appearance of the ablated larvae at 7 dpf, with further incubation in a controlled nursery with regular feeding we observed a steep decline in survival beginning at approximately 11 dpf, with only 25% of the MTZ-treated NTR animals still surviving at 23 dpf, compared to a much lower loss of animals in all of the control groups (Log-rank Mantel-Cox test, *p*-value=<0.0001) (**Fig. 8L**).

Ablation of LBCs in adult zebrafish is also lethal. When adult NTR zebrafish were treated with 10 mM MTZ for 24 hours then rinsed and maintained in E3 media without drug (**Fig. 8M**), 62.5% of animals died by the end of the 24 hour MTZ treatment (n=10/16), and all remaining MTZ-treated animals (n=6/16) died within the following 24 hours (**Fig. 8N**). During the first treatment day MTZ-treated animals exhibited marked changes in behavior including reduced swimming and reduced responsiveness to stimuli (**Supp. Movie 5**). Upon closer examination, MTZ-treated animals display global cephalic hemorrhage, with blood pooling in areas surrounding the brain (arrows) such as the olfactory bulbs, optic tecta, and cerebellum (**Fig. 8P,R**). Control NTR animals that were not treated with MTZ (n=16/16) showed 100% survival and displayed no signs of any behavioral or hemorrhage phenotypes (**Fig. 8O,Q, Supp. Movie 5**). MTZ treatment of NTR fish resulted in rapid ablation of LBCs, but it did not result in the immediate loss of FGPs (**Fig. 8S-V**). 12 hours after MTZ treatment FGPs were still present, although they appeared less spindle-like in shape, had many fewer filopodial projections, and showed signs of rounding and blebbing suggestive of dying cells (**Fig. 8T,V**). As noted above, FGPs are known for their robust capacity to internalize materials from the external environment, and the FGPs in MTZ-treated NTR adults were filled with large mCherry-labeled internal vesicles (**Fig. 8T,V, Supp. Figure 8 and Supp. Movie 6**), which were completely absent in control untreated NTR animals (**Fig. 8S,U, Supp. Figure 8**), indicating that FGPs were actively taking up debris from dying LBCs.

## DISCUSSION

We have carried out a comprehensive anatomical, cellular, and molecular analysis of the zebrafish meninges and its constituent cell populations. Histological and ultrastructural characterization shows that the adult zebrafish cephalic meninges are a complex multilayered tissue with a mammalian-like structure, including a bilayered skull-associated dural meninges and a compound brain-associated leptomeninges, contrary to some classical reports describing a single-layered “primitive meninx” in several different fish species (eg., (Sagemehl, 1884; Sterzi, 1902). Using single cell RNA sequencing (scRNA-seq), we identified distinct populations of cells in the dural meninges and leptomeninges. Both layers have blood vessels and abundant immune cell populations, particularly the dural meninges. As in mammals, lymphatic vessels are found only in the skull-associated dural meninges, although the brain-associated leptomeninges includes a population of perivascular scavenger cells (FGPs, aka muLECS or BLECs) with lymphatic-like molecular identity that emerge from primitive lympho-venous endothelium (Bower et al., 2017; van Lessen et al., 2017; Venero Galanternik et al., 2017). We have also identified and characterized a second population of cells unique to the leptomeninges, ependymin-expressing Leptomeningeal Barrier Cells or “LBCs.” Although LBCs are molecularly similar to and share a close lineage relationship with meningeal fibroblasts, which have also been reported in mammals (DeSisto et al., 2020), they represent a separate and distinct cell population. We show that LBCs are the primary epithelial cell type of the leptomeninges, within which blood vessels, FGPs, and other cell types are embedded, and we demonstrate that the presence of LBCs is essential for juvenile zebrafish survival and that acute ablation of LBCs in adult zebrafish is lethal. Together, our findings establish the zebrafish as an important new model for experimental and genetic analysis of the vertebrate meninges.

The mammalian meninges are a complex set of tissues composed of three main layers, the *dura mater, arachnoid mater,* and *pia mater* (Greenberg et al., 1994; O’Rahilly and Muller, 1986). The zebrafish meninges have an equally complex, mammalian-like structure. The dural meninges or *dura mater* is the most external meningeal layer, found immediately below the skull. Like mammals, zebrafish have a bilayered dural meninges pachymeninges) that includes a periosteal dura immediately juxtaposed to the skull as well as an adjacent deeper meningeal dura. In addition to blood vessels, the zebrafish dural meninges contain a network of lymphatic vessels (Castranova et al., 2021). The dural meningeal lymphatic plexus has functional *mrc1a+, lyve1b+, flt4+, stab1/stab2+* lymphatic vessels that take up interstitial fluid, transport immune cells, and respond to *vegfc*-induction (Castranova et al., 2021), and they represent a homologous lymphatic network to the mammalian meningeal lymphatics recently described in mice and humans (Aspelund et al., 2015; Louveau et al., 2015). The zebrafish dural meninges are also rich in immune cells. As noted above, immune cell clusters make up 34.2% of the cells captured in our dural scRNA-seq samples. After erythrocytes, T cells represent the largest cell cluster in our dural scRNA-seq data set, and patrolling T cells are readily observed in live imaging of the zebrafish dural meninges. B cell, neutrophil, and macrophage clusters are also clearly identified. Epithelial and ciliated cell clusters were also identified in the dural scRNA-seq that likely include cells that make up the epithelial lining of the dural layers, although the specific localization of these cell populations was not explored in detail. Pigment-containing cells are also present in the zebrafish dura mater, including melanin-rich cells, iridophores and xanthophores. Melanocytes have been described in the meninges of mice, rats, cats and humans (Fetissov et al., 1999; Goldgeier et al., 1984; Morse and Cova, 1984), although there appear to be fewer meningeal melanocytes in these animals than in fish and they are generally found localized to fewer areas within the meninges. The role of meningeal melanocytes in animals with thick, opaque skulls remains unclear, however there are suggestions that these cells may play a role in modulating immune and/or vascular responses (Miniati et al., 2014; Tsatmali et al., 2002). The larger numbers of pigment containing cells found in the zebrafish meninges may reflect the need for additional UV protection for the meninges and brain given the thin translucent skulls of zebrafish and their natural habitat in sunny, shallow waters of South Asia (Parichy, 2015). It would be interesting to compare the meningeal pigment cells in zebrafish with those of other fish adapted naturally to dark environments.

Like mammals, zebrafish also have a separate leptomeninges in addition to the dural meninges. In humans the leptomeninges is divided into distinct *arachnoid mater* and *pia mater* layers. In zebrafish these layers are not clearly separated, but transmission electron microscopy reveals the presence of a multilayered leptomeninges separated from the brain parenchyma by a well-defined basal lamina and adjacent glia limitans. Like its mammalian counterpart, the zebrafish leptomeninges contains blood vessels but no lymphatic vessels. Instead, the zebrafish leptomeninges contains lymphatic-like perivascular scavenger cells (FGPs/muLECS/BLECs) that actively take up materials from the cerebrospinal fluid and interstitial spaces and incorporate them into large intracellular vesicles (Bower et al., 2017; van Lessen et al., 2017; Venero Galanternik et al., 2017). Although FGPs arise from pre-existing primitive veins and express many if not most lymphatic markers, they do not assemble into tubular structures. We confirmed that these cells are confined to the leptomeninges and are not present in the dural meninges. Based on their location, scavenger behavior and gene expression, FGPs are most similar to mammalian meningeal perivascular macrophages (PVMs) or border-associated macrophages (BAMs), cells that reside at the interface between the brain and peripheral structures such as the meninges and choroid plexus (Da Mesquita and Rua, 2024; Mendes-Jorge et al., 2009; Sun and Jiang, 2024; Wen et al., 2024; Zheng et al., 2022), although the similarities and differences between zebrafish FGPs and mammalian PVMs or BAMs need further investigation.

In addition to FGPs, our scRNA-seq revealed a separate novel leptomeningeal cell population. We identified two fibroblast-like cell clusters in our zebrafish leptomeningeal scRNA-seq data set, a smaller cluster corresponding to mammalian meningeal fibroblasts (DeSisto et al., 2020; Pietila et al., 2023; Siegenthaler and Pleasure, 2011) and a separate, larger cluster containing cells strongly expressing ependymin, a cerebrospinal fluid glycoprotein previously implicated in dominant behavior, neuro-regeneration and memory establishment in teleosts (Hoffmann, 1994; Hoffmann and Schwarz, 1996; Sneddon et al., 2011; Suarez-Castillo et al., 2004). Although these two cell types represent distinct mesenchyme-derived cells, they share a close lineage relationship during early zebrafish larval development and share common expression of a number of genes. Using new transgenic lines we generated we showed that ependymin-expressing cells are unique to the leptomeninges and that they appear to make up the majority of the epithelial structure of this meningeal layer. We have designated these new cells Leptomeningeal Barrier Cells, or LBCs. Using genetic ablation of LBCs via transgenic Nitroreductase-driven conversion of Metronidazole, we show that ablation of LBCs during early larval development subsequently results in progressive death in early juvenile animals. The requirement for an intact leptomeninges and the precise reason for death in juveniles will need further investigation. Acute ablation in adult zebrafish using similar methods results in death within one day, with evidence of extensive intracranial hemorrhage. Although the complete ablation we tested here is lethal, limited partial inducible ablation of LBCs in adult zebrafish is likely survivable, and may serve as a valuable new model for meningeal traumatic brain injury in the future. In mammals, epithelial-like meningeal arachnoid barrier cells (“AB cells”) have been also shown to be critical for the barrier function of the meninges. Like LBCs, AB cells originate from fibroblastic mesenchymal precursor cells during early embryonic development but then begin to express epithelial and tight junction proteins and take on an epithelial character (DeSisto et al., 2020; Uchida et al., 2019), with recent work showing that downregulation of Wnt signaling is needed for this transition (Derk et al., 2023). Although *ependymin* itself is a teleost-specific gene, mammals have *ependymin*-related proteins (EPDRs, also called “Mammalian Ependymin-related proteins” or “MERPs” in mammals) that are likewise found in the extracellular matrix of the central nervous system, including the meninges. It will be interesting to investigate whether AB cells or similar fibroblast-like epithelial cells in the mammalian leptomeninges express high levels of MERPs, and if they represent direct or only functional orthologs of LBCs.

Our findings establish the zebrafish as an important new model for experimental and genetic analysis of the vertebrate meninges, with a mammalian-like multilayered meninges located under a thin translucent skull that makes it possible to carry out live imaging of the meninges in intact, living animals. Despite their relatively superficial location, the mammalian meninges are generally not accessible in intact animals as they are covered by hair, skin, and a thick bony skull that must be manipulated to observe or experimentally handle the meninges (Coles et al., 2017b; Cramer et al., 2021; Dorand et al., 2014). Hence, with relatively few exceptions, most mammalian studies on meningeal development and cellular composition rely on use of sectioned post-mortem tissues rather than live *in vivo* analyses. Live imaging of the murine meninges requires invasive procedures to generate “windows” for imaging that themselves result in extensive injury (Coles et al., 2017a). The zebrafish provides a useful alternative. Zebrafish embryos and larvae are transparent, and in adults the brain surface is readily observed through the thin translucent skulls of intact, unmanipulated animals raised on pigment-deficient backgrounds. Methods have been developed for long-term time-lapse confocal imaging of intubated adult zebrafish and they are regularly used for live *in vivo* imaging studies (Castranova et al., 2022; Chow et al., 2020; Greenspan et al., 2024; Xu et al., 2015). Adult fish are also amenable to genetic, chemical, and other experimental manipulations, further expanding their usefulness as a model organism for meningeal studies. In the future, development of new cerebrovascular/meningeal injury or infection tools and methods for the zebrafish will help make the zebrafish a particularly useful model for studying and developing new therapeutic approaches for treating pathologies affecting the meninges and brain such as traumatic brain injury (TBI), cerebrovascular injury (CVI), neuroinflammation, and meningitis.

## MATERIALS AND METHODS

### Fish Husbandry and Fish Strains

Fish were housed at the National Institutes of Health zebrafish dedicated recirculating aquaculture facility (4 separate 22,000L systems) in 6L and 1.8L tanks. Fry were fed rotifers and adults were fed Gemma Micro 300 (Skretting) once per day. Water quality parameters were routinely measured, and appropriate measures were taken to maintain water quality stability (water quality data available upon request). The following transgenic fish lines were used for this study: *Tg(mrc1a:eGFP*)*^y251^* (Jung et al., 2017), *Tg(−5.2lyve1b:DsRed)^nz101^*(Okuda et al., 2012), *Tg(kdrl:mcherry)^y206^* (Gore et al., 2011), *Tg(lyz:dsred2)^nz50^*(Hall et al., 2007), *Tg(lck:mcherry)^ns107^* (Amanda et al., 2022), *Tg(lck:egfp)^cz1^* (Langenau et al., 2004), *Tg(cd79b:egfp)^fcc89^*(Liu et al., 2017), *Tg(mpeg1:egfp)^gl22^* (Ellett et al., 2011), *Tg(uas:ntr-mcherry)^c264^* (Gutnick et al., 2011) and *TgBAC(pdgfrb:EGFP)^ncv22^*(Ando et al., 2016). As part of this study we also generated the following new lines for use: *Tg(epd:mcherry)^y715^, Tg(epd:gfp-caax)^y716^*and *Tg(epd:gal4-VP16)^y717^* using the Tol2/Gateway system (Kwan et al., 2007). Pigmented animals were maintained in the EK wild type background and most of the lines imaged were maintained and imaged in a *casper* (*roy, nacre*) double mutant genetic background (White et al., 2008) in order to increase clarity for visualization of the meninges by eliminating melanocyte and iridophore cell populations from the top of the head.

### Image Acquisition

Confocal images of intracranial lymphatics were acquired using a Nikon Ti2 inverted microscope with Yokogawa CSU-W1 spinning disk confocal, Hamamatsu Orca Flash 4 v3 camera, and a Nikon AX-R 2K large field of view hybrid scanning inverted confocal microscope. Objectives used where the (i) Nikon Apochromat lambda D 4X Air 0.2 N.A., 10X Air 0.45 N.A., 20X air N.A. 0.80, the (ii) Nikon APO Lambda S 20X WI water immersion N.A. 0.95, 25X silicone immersion N.A. 1.05, 40X WI 1.15 N.A., and the Apochromat 60X OIL TIRF NA 1.49 immersion. The large size of the juvenile and adult zebrafish heads often required tile acquisitions that were later stitched using Nikon Elements software. Stereo microscope pictures of adult or embryonic animals were taken using a Leica M205 microscope or a Leica Ivesta 3 microscope with integrated digital color camera using MultiFocus focus stacking.

### Transmission Electron Microscopy

Wild type zebrafish were euthanized in an ice bath and fixed overnight at 4°C in 4% PFA with 2.5% glutaraldehyde, made in 0.1M sodium cacodylate buffer, pH 7.4. The tissue samples were then rinsed in 0.1M sodium cacodylate buffer, and post-fixed in 1.0% osmium tetroxide in 0.1M sodium cacodylate buffer for 60 minutes at room temperature. Next, the samples were rinsed in MilliQ water, then incubated in 3% uranyl acetate (aqueous) for 60 minutes at room temperature, followed by several MilliQ water rinses. Samples were dehydrated in 50%, 70%, and 90% EtOH 2 × 10 mins each, then three 100% ETOH exchanges (10 mins each), followed by three 100% propylene oxide exchanges (10 mins each). A 1:1 mixture of propylene oxide and Embed-812 resin (Electron Microscopy Sciences, Hatfield, PA) was allowed to infiltrate the samples for 2 days. The samples were then transferred to 70% Embed-812 resin/30% propylene oxide for 2 hours, followed by five exchanges in 100% fresh Embed-812 resin for 1 hour each. Samples were then embedded in fresh Embed-812 resin and polymerized in a vacuum oven set at 60°C for 24 hours. Thin sections were cut on a Leica EM-UC7 Ultramicrotome (90 nm). Thin sections were picked up and placed on 200 mesh copper grids and post-stained with lead citrate. Imaging was accomplished using a JEOL-1400 Transmission Electron Microscope operating at 80kV and an AMT BioSprint 29 camera.

### Confocal Image Processing

Confocal images were processed using the Nikon Elements software, ImageJ, Adobe Photoshop 2023 and Imaris. All images where mCherry or dsRed2 were used, are displayed in magenta when in combination with GFP as suggested by color-blind palettes. Unless otherwise specified, maximum intensity projections of confocal stacks are shown. Focus stacking of confocal images with DIC was done using Nikon Elements EDF (Extended Depth of Focus). 3D rotation movies and time-lapse movies were made using Nikon Elements and exported to Adobe Premiere Pro CC 2019. Adobe Premiere Pro CC 2019 and Adobe Photoshop CC 2019 were used to add labels and arrows to movies and to add coloring or pseudo-coloring.

### Tissue Sectioning and H&E Staining

Adult zebrafish were euthanized on an ice bath and their brains were dissected out manually and washed in 1X PBS, without calcium or magnesium. For H&E staining, brains (n = 3) were embedded in O.C.T Compound media (Tissue-Tek) and sectioned into 10 μm thick sections (Histoserv, Inc.). Adult mice (n = 3) were anesthetized with Isoflurane and perfused intracardially with 50 mL of 1X PBS, followed by 50 mL of 4% PFA. The brains were manually dissected and place in 1X PBS for a wash, followed by a gradient of 5%, 10% and 20% Sucrose/PBS washed with 0.01% Sodium Azide. Brains were then cryo-sectioned into 50 μm slices and stained with H&E staining (Histoserv., Inc.).

### Adult Tissue Dissections

Animals were euthanized on an ice bath for 5 to 10 minutes. Brains and skulls were manually dissected by placing the animals on a 150 x 20 mm silicone pad dissecting dish (VWR #100491-622). Dorsal portions of skull caps covering the optic tectum, cerebellum and telencephalon were manually removed using Dumont Tweezers #55 (VWR #72707-01). Exposed brains were carefully removed from the calvaria by severing the brain-spinal cord connection and nicking the optic nerves connecting the brain to the eyes. Dissected skulls and brains were immediately rinsed in 1X PBS, no calcium, no magnesium (Quality Biological #114-340-131) to remove any remnants of blood, fat or bones. For imaging purposes, dissected brains and skulls with intact meninges were immediately used for confocal imaging, electron microscopy and cell dissociations.

### Preparation of Meningeal Cell Suspensions for scRNA-seq

A total of 20 adult zebrafish were used to prepare leptomeningeal cell suspensions for scRNA-seq (10 females, 10 males). Animals were euthanized in an ice water bath one at a time and brains were carefully dissected, preserving the optic tectum intact. Leptomeningeal linings were manually removed from the optic tectum and placed in 1X PBS with no calcium or magnesium (Quality Biological Cat# 114-340-131). This tissue was transferred to a freshly prepared papain solution, following manufacturing instructions (Worthington Biochem Cat #LK003150) with some modifications as suggested by previous zebrafish brain isolates used for scRNA-seq (Raj et al., 2018). Briefly, dissected leptomeningeal tissues were incubated at 34°C for 20-30 min in 2 mL of 10 units/papain in neurobasal media (Life Technologies Cat # 21103049) equilibrated with 95% O_2_:5% CO_2_ as described in (Raj et al., 2018) with gentle interval pipetting using a p1000 and then p200 filtered tips. Dissociated cells were then filtered through a 40 mm cell strainer (Gen Clone Cat #25-375). Supernatant was discarded and cells were spun for 4 min at 500 g. The cell pellet was resuspended in 1mL of fresh oxygenated Ovomucoid/DNase inhibitor solution prepared as described by the manufacturer (Worthington Biochem Cat #LK003150), spun at 500 g for 4 minutes, and the pellet was rinsed twice with DPBS media with no calcium, no magnesium (Thermo Fisher #14190144, spinning the sample in between rinses. The final pellet was resuspended in 500 mL of DPBS and checked for cell viability.

A total of 25 adult zebrafish were used to prepare dural meningeal cell suspensions for scRNA-seq (12 females, 13 males). Skull caps were carefully dissected from animals euthanized as above and each was placed dorsal side down into the internal well on the inside of a 0.2 ml PCR flat tube cap that had been previously filled with 3% agarose. Once all skulls were collected and placed in their own PCR tube cap, 10 μL of freshly made papain solution (as described above) was added to the inner ventral side of each cap and then incubated for up to 30 minutes, gently pipetting up and down every other minute, until all observable meningeal tissue was detached from the inside of the skull. The dissociated dural cells were then processed similarly to the leptomeningeal samples described above.

All cell suspensions were assessed for viability and numbers on a LUNA-FL (Logos Biosystems) and diluted to the optimum concentration of 700 cells/ml using DPBS. 10,000 cells were loaded onto the 10x Genomics ChromiumX controller. A total of four dural suspensions (DM1-4) and six leptomeningeal (LM1-6) were used.

### Single Cell Data Analysis

Alignment of sequencing reads and processing into a digital gene expression matrix was performed using Cell Ranger. DM1,2 and LP2,3 – v6.0.1, DM3,4 – v7.0.0, DM5,6 and LP4,5 – v7.0.1. Reads were aligned against GRCz11 using gene annotations from ENSEMBL release 99. Cells were processed and analyzed using R version 4.1.2 and Seurat version 4.0.6 (https://doi.org/10.1016/j.cell.2019.05.031), for the code used or this please refer to https://github.com/nichd-Weinstein/Meninges. Doublets called by scDblFinder (v1.8.0) in each of the six pachymeninges (DM) datasets, and in each of the four leptomeninges (LP) datasets, were removed. Cells with abnormally high (> 5000) or low (< 200) numbers of detected features, or with abnormally high mitochondrial content (> 10%) were removed. The remaining cells were processed in Seurat with default settings, including integration of the six pachymeninges datasets, and separately, the integration of the four leptomeninges datasets.

Following initial clustering of the pachymeninges data (Leiden clustering at the default resolution of 0.8), it was noted that many cells in the (integrated) pachymeningeal dataset expressed elevated mitochondrial and ribosomal genes (**Supp. Fig 1**). These cells are represented in **Supp. Fig. 1B** as clusters 1-6, 8, 9, 11 and 12. Cells in these clusters (a total of 8,346 cells) were removed and retained cells were reprocessed from the beginning, including integration and clustering. Further manual annotation was carried out to reduce six erythrocyte clusters into one, four T cell clusters into one, to split macrophages and myeloid precursors into separate clusters, and to divide endothelial and mural cells into separate clusters (see **Supp. Table 1** for a list of genes used for manual annotation of cell identities, https://zenodo.org/records/15178518).

Following initial clustering of the leptomeninges data (Leiden clustering at the default resolution of 0.8), and the subsequent removal of cells with elevated mitochondrial and ribosomal genes, a significant number of cell clusters corresponding to mature neuronal populations (a total of 17,109 cells) were identified by their expression of GABA and glutamatergic genes as shown in **Supp. Fig. 2** and **Supp. Tables 1, 2**. Since these were the result of cross-contamination from closely attached brain tissue during manual dissections, clusters containing these cells (shown in **Supp. Fig 2B-D** as Cluster 1,2, 4-6, 8-10, 12, 14-18, 28 and 33) were filtered out and the retained cells reprocessed from the beginning including integration and clustering. Since the pachymeninges is not directly attached to the brain and remains connected to the skull upon dissection, neuronal contamination of that dataset was minimal, and neuronal filtering was not needed. Further manual annotation of leptomeninges clustering was carried out to reduce six erythrocyte clusters into one, four residual unfiltered glial cell clusters into one, and to separate out a small B cell cluster (identified by expression of *igic1s1*), using expression of the genes listed in **Supp. Table 2** as a guide. The resulting pachymeningeal and leptomeningeal scRNA-seq datasets shown in **Fig. 3B-C** were used for all subsequent analyses.

Genes were considered differentially expressed if the adjusted P value was lower than 0.01. A table of the resulting genes with the highest expression values in each cluster can be found in **Supp. Table 1**. As noted above, each of the clusters in **Figure 3B,C** was manually annotated based on an extensive survey of well-known tissue- and cell type–specific markers. These markers were identified through a variety of databases (Human Protein Atlas, The Zebrafish Information Network, and Daniocell) and an extensive literature search covering mammalian and piscine publications. For each cell type, at least four marker genes were identified, listed in **Supp. Table 2**.

### Inference of developmental trajectories using URD

For URD pseudotime analysis (Farrell 2018), the Daniocell mese.21, mese.29, and mese.32 clusters were used as input. Some of the cells forming a bridge between mese.29 and mese.32 were removed from the analysis. 24–34 hpf cells from cluster mese.21 were used as the root, and 120 hpf cells from clusters mese.29 and mese.32 were defined as two tips. A diffusion map was calculated using the function URD::calcDM with parameters nn = 100 and sigma = 7.5. Next, pseudotime was computed using the function URD::floodPseudotime (n = 100, minimum.cells.flooded = 2). Then biased random walks were simulated starting at each terminal using the function URD::simulateRandomWalkfromTips and a branchpoint was called using the following parameters – optimal.cells.forward = 10, max.cells.back = 20, n.per.tip = 25000, root.visits = 1, max.steps = 5000. Next, a branching tree structure was fit to the trajectory using the URD::buildTree command with the following parameters: divergence method: “preference”, cells.per.pseudotime.bin = 25, bins.per.pseudotime.window = 5, p.thresh = 0.1). Using the URD branching tree as a framework, the URD:aucprTestAlongTree() function was used to find differentially expressed genes of each lineage using the parameters: must.beat.sibs = 0.3, auc.factor = 0.6, log.effect.size = 0.4, max.auc.threshold = 0.85. Genes were called differentially expressed if they were expressed in at least 10% of cells in the branch under consideration and were 0.6 times better than a random classifier for the population as determined by the area under a precision-recall curve. Branchpoint plots of selected differentially expressed TFs were created using the URD::branchpointPreferenceLayout command to calculate a preference layout for the branchpoint based on the visit frequency by random walks from the two segments, that was used as an input to plotting branchpoint plots using the plotBranchpoint function (x-axis: populations = “mFB” and “LM”; y-axis: pseudotime = “pseudotime”.

### Identifying gene cascades along developmental trajectories

To construct gene expression cascades along the two branches, gene expression dynamics within leptomeninges and meningeal fibroblasts were fit using smoothed spline curves using the URD::geneSmoothFit() and parameters – EPDs: moving.window = 1, cells.per.window = 5, spar = 0.9, and meningeal fibroblasts: moving.window = 4, cells.per.window = 8, spar = 0.9. For this trajectory, genes were further compared between the two branches to select genes that are specific to one branch or another or markers of both. This strategy was used to differentiate between meningeal precursors, EPDs, and fibroblasts. Pseudotime was then cropped at the branchpoint of these two cell types and the smoothed spline curves were plotted using the function URD::plotSmoothFitMultiCascade.

### *ependymin*-dependent transgenic line generation

Using the Tol2/Gateway system (Kwan et al., 2007), we generated three stable transgenic lines for this study; *Tg(epd:mcherry)^y715^, Tg(epd:gfp-caax)^y716^* and *Tg(epd:gal4-VP16)^y717^*. Briefly, using a BAC clone (CH211-69l14) as template, we PCR cloned a 4.9 Kb sequence upstream or the ATG site of the zebrafish *ependymin* gene (*Epd-F*:GGGGACAACTTTGTATAGAAAAGTTGTATAACTTATTTACACTATGGGGTGTTTTT and *Epd-R*:GGGGACTGCTTTTTTGTACAAACTTGATTCAGACTCTGCCTTTTATACATAACC). This upstream sequence was selected based on conserved sequence comparison to other teleosts, the presence of CpG islands, a TATA box sequence and an overlapping 2.0 Kb previously published minimal regulatory region able to drive the expression of lacZ (Rinder et al., 1992). Upon Sanger sequencing confirmation of the desired promoter region, we used the Tol2kit/Gateway system to insert the sequence into a p5E entry vector and used the Tol2Kit pME clones pME-EGFP-CAAX, pME-mCherry and pME-Gal4VP16, and p3E-polyA to assemble the final constructs in a pDestTol2pA vector (Kwan et al., 2007). Transgene constructs were injected into 1-cell stage wild type zebrafish embryos together with with Tol2 transposase to ensure random genomic insertion. F0 (injected) embryos showing fluorescent expression were selected and raised to adulthood, then outcrossed individually to EK wild type animals to test for germ line transmission. Transgene-expressing F1 progeny were raised to adulthood and used to generate stable F2 families from which these lines have been maintained.

### Confocal imaging of Dissected Adult Brains

Freshly dissected adult transgenic brains were rinsed in 1X PBS, with no calcium and no magnesium and immediately mounted on a drop of 1% Methylcellulose in E3 solution, with their dorsal side facing down on a 35 mm imaging dish (MatTek # P35G-1.5-14-C). The brains then were covered with a cover slip to avoid drying during image acquisition and immediately imaged on an inverted confocal microscope as described above.

### Live Confocal imaging of Zebrafish Larvae, Juveniles, and Adults

Living transgenic larvae between 2-7 days post fertilization (dpf) were anesthetized in E3 media containing 4mg/mL MS222/Tricaine (Thermo Fisher # NC0872873) and mounted in 0.8% Low Melting Agarose (LMP) in E3 media on a 35 mm imaging dish (MatTek # P35G-1.5-14-C) either laterally or with dorsal side against the glass. Once the LMP agarose had solidified, the embryos were covered with E3 media containing tricaine and placed on an inverted confocal for imaging acquisition. For long term imaging, mounted embryos were placed on a confocal stage heating chamber and kept at 28.5°C during acquisition period to ensure proper development.

Adult and juvenile fish were mounted by anesthetizing them in 126-168 mg/L tricaine in system water and then placed into a slit in a sponge (Jaece Identi-Plugs L800-D) moistened with tricaine water, cut into a rectangle to fit inside a single chamber imaging dish (Lab-TekII #155360). The sponge containing the fish was placed into the imaging dish with the fish’s head against the glass (fish upside down) and the chamber was filled with tricaine water. Upon imaging completion, the animals were revived by placing them in a tank of fresh system water. If no signs of distress were observed, the animals were placed back into a recirculating system. If signs of stress were observed, or the animals were not able to recover cognition and swimming capacities, the animals were immediately euthanized on an ice bath.

### Angiography Injections

Injections were performed as described by Yaniv et al. (Yaniv et al., 2006). All injections were performed using a Drummond Nanoject II microinjector (Item# 3-000-204) with pulled glass capillary needles (Drummond item # 3-00-203-G/X). One or two injection boluses were given at each injection site with a volume setting of 50 nL. Angiography assays were done using DRAQ5 (5 mM) injected into the caudal axial vasculature (Thermo Scientific Product # 62254).

### Embryonic and adult *ependymin*-expressing cell ablations

For sustained LBC ablations in larval zebrafish, incrossed embryos were raised to 5 dpf and sorted for *Tg(epd:gal4-vp16);Tg(uas:ntr-mCherry);Tg(mrc1a:eGFP)*, or only *Tg(mrc1a:eGFP).* Both groups were placed in either 1% DMSO E3 media or 10 mM Metronidazole, 1% DMSO (Thermo Fisher # H60258.14) solution in E3 at a density of 5 embryos/mL. Fish were treated from 5 hpf to 7 dpf and viability was assessed on each day of treatment, with daily changes of media. A sample of ∼10-15 embryos per treatment were mounted in 1% LMP agarose and imaged on an inverted confocal microscope.

For acute LBC ablation in adult zebrafish, 6-month-old *Tg(epd:gal4-vp16);Tg(uas:ntr-mCherry);Tg(mrc1a:eGFP)* transgenic *casper* and *nacre* zebrafish were used. Animals were placed on system water containing either 2% DMSO (n=16, 1:1 Female to male ratio) or 10 mM Metronidazole (Thermo Fisher # H60258.14, n=16, 1:1 Female to male ratio) for up to 24 hours. Following established IACUC protocols, animals with LBC ablations were quickly i) imaged on a Leica Ivesta 3 dissecting microscope and euthanized immediately with the purpose of imaging their freshly dissecting brains, or ii) used to live image their meninges through the intact skull.

## ACKNOWLEDGMENTS

We thank all the Weinstein Lab members for their suggestions and valuable discussions. We are grateful to Drs. Harold Burgess and Richard Dorsky for their mentoring to MVG, and Dr. Amber Stratman for thoughtful comments on the manuscript.

## SUPPLEMENTAL FIGURE LEGENDS

**Supplemental Figure 1.**
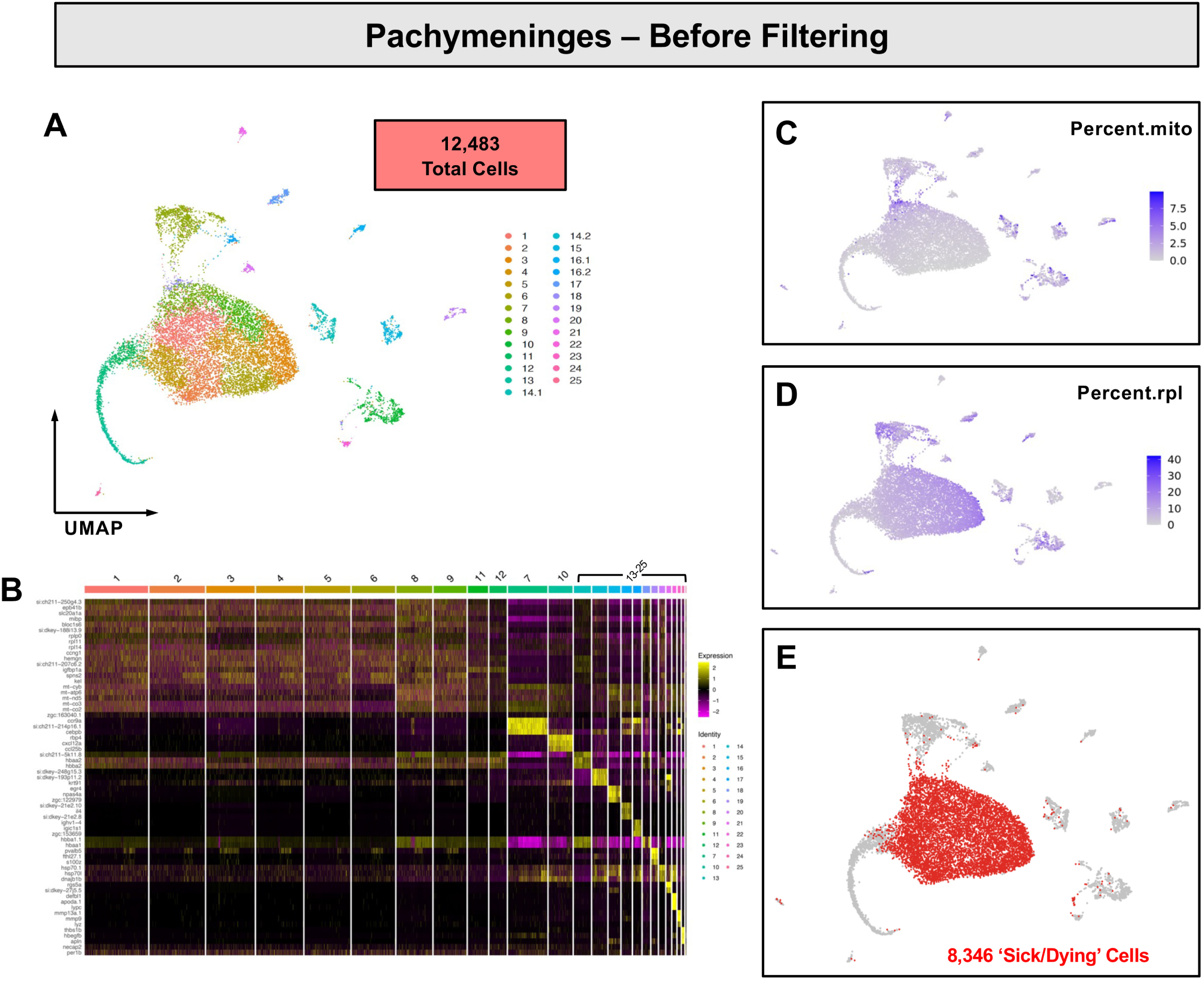
Filtering dying cells from the pachymeningeal (dural meningeal) scRNA-seq data set. **A,** Uniform Manifold Approximation and Projection (UMAP) plots of unfiltered single cell RNA-seq (scRNA-seq) data obtained for dissected pachymeninges (dural meninges), including a total of 12,483 cells. **B,** Heat map of genes enriched in pachymeninges scRNA-seq cell clusters. Clusters 1-12 show similar expression of mitochondrial and ribosomal genes commonly enriched in stressed and/or dying cells. **C,D,** Feature plots of pachymeninges scRNA-seq data showing cells with high levels of mitochondrial (C) or ribosomal (D) gene expression. **E,** Feature plot highlighting cells that showed enriched mitochondrial and ribosomal gene expression and limited cluster-specific gene expression (red). All of these cell clusters (8,346 cells in total) were removed from the final filtered pachymeningeal UMAP showed in Figure 3B.

**Supplemental Figure 2.**
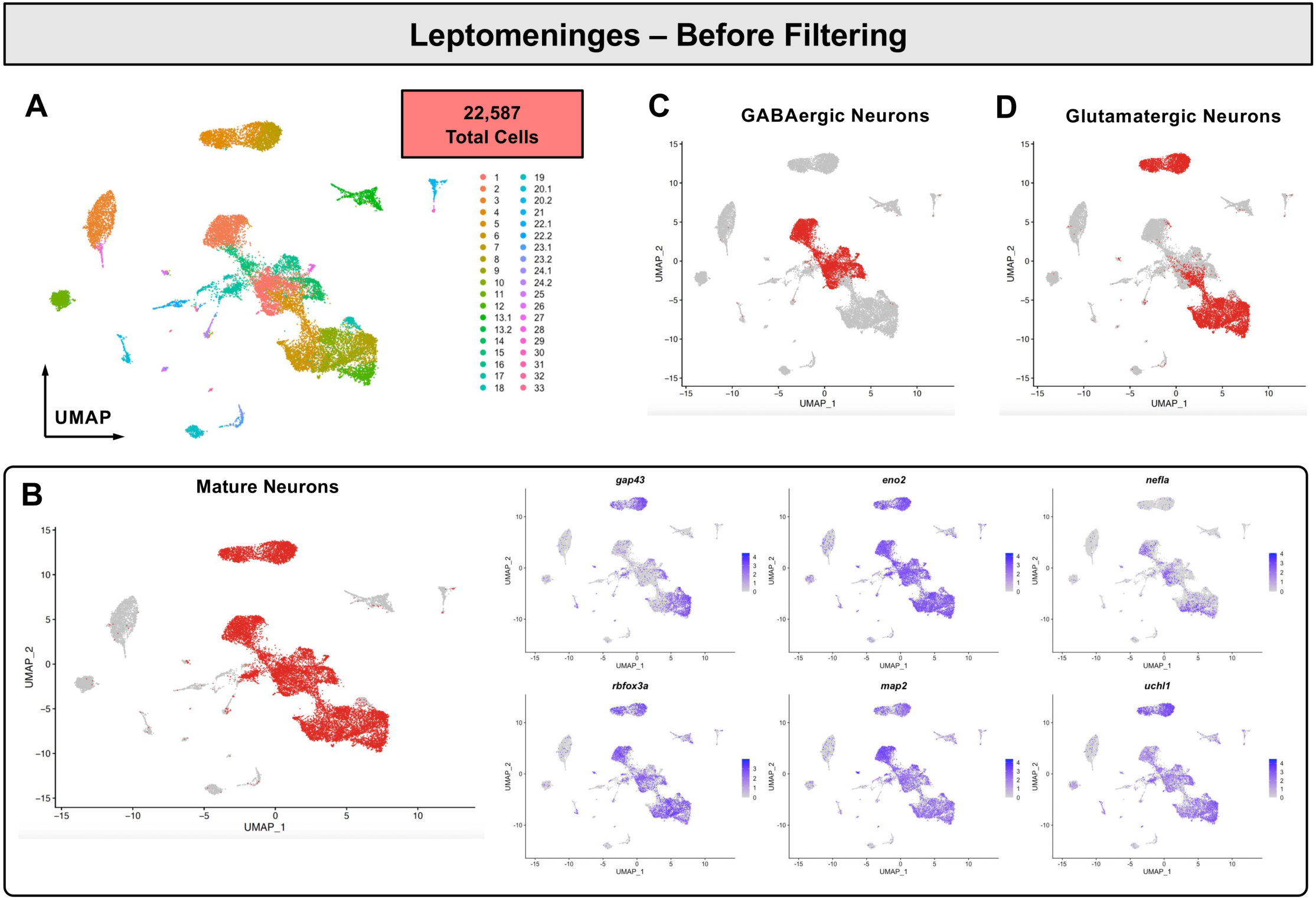
Filtering neural cells from the leptomeningeal scRNA-seq data set. **A,** UMAP plot of unfiltered single cell RNA-seq (scRNA-seq) data obtained for dissected leptomeninges, including a total of 22,587 cells. **B,** Feature plot showing leptomeningeal scRNA-seq cell clusters with a mature or precursor neuronal cell identity based on their gene expression profile (red). Individual feature plots show the distribution of mature neuron markers expressed by these cells (*gap43, eno2, map2, nefla, uchl1, rbfox3a*). All of these cell clusters 17,109 cells in total) were removed from the final filtered leptomeningeal UMAP showed in Figure 3C. **C,D,** Feature plots highlighting cells in red expressing markers consistent with a GABAergic (C) or glutamatergic (D) neuronal identity.

**Supplemental Figure 3.**
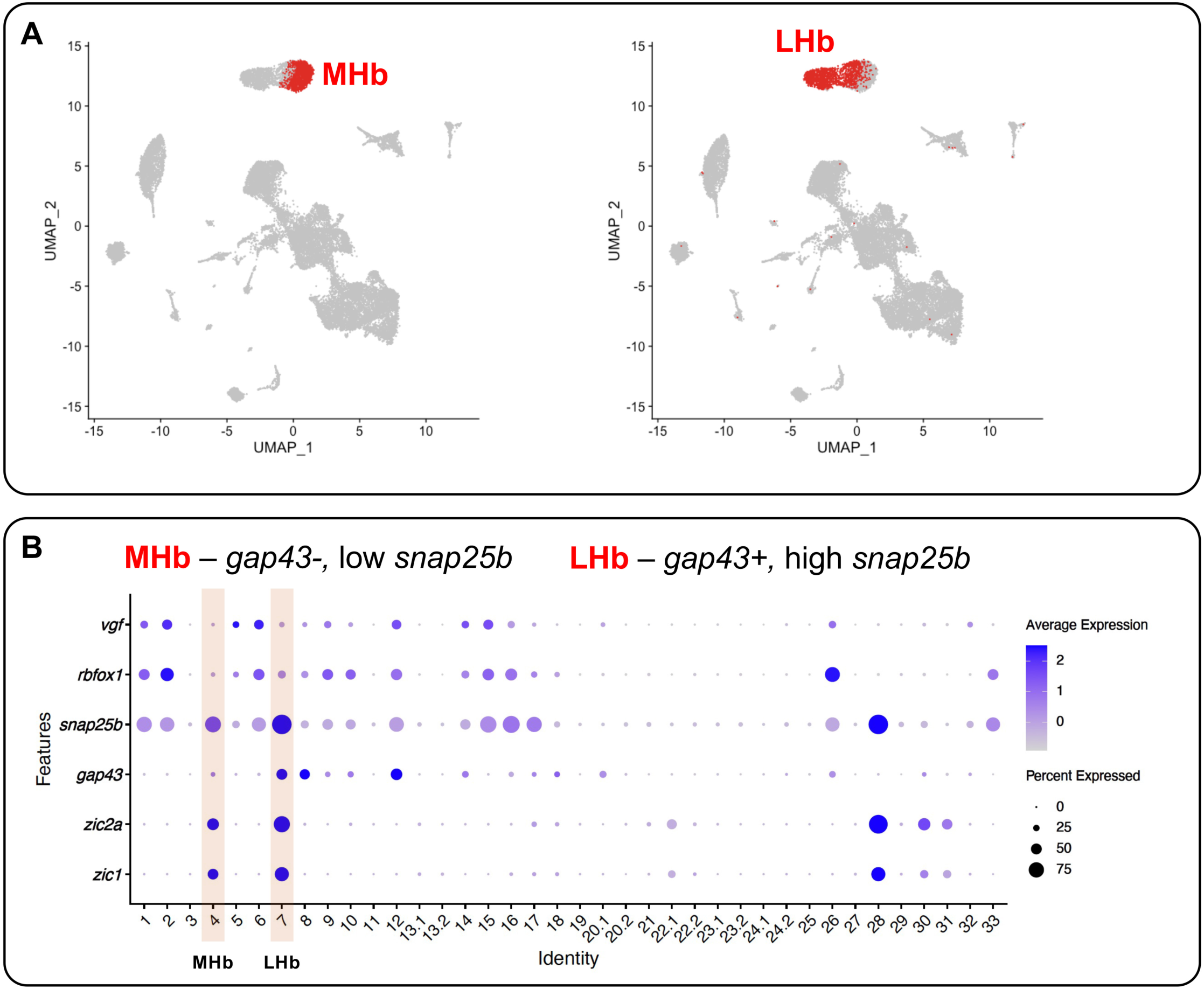
Habenular neurons in the leptomeningeal scRNA-seq data set. **A,** Feature plots highlighting leptomeningeal scRNA-seq cell clusters (red) containing cells with medial habenula (MHb, left) or lateral habenula (LHb, right) identity. **B,** Dot plot showing genes expressed by leptomeningeal scRNA-seq cell clusters including putative MHb and LHb clusters (highlighted by light red bars). The MHb shows almost no expression of *gap43* and lower expression of *snap25b*, while the LHb shows characteristic high expression of *gap43* and *snap25b*.

**Supplemental Figure 4.**
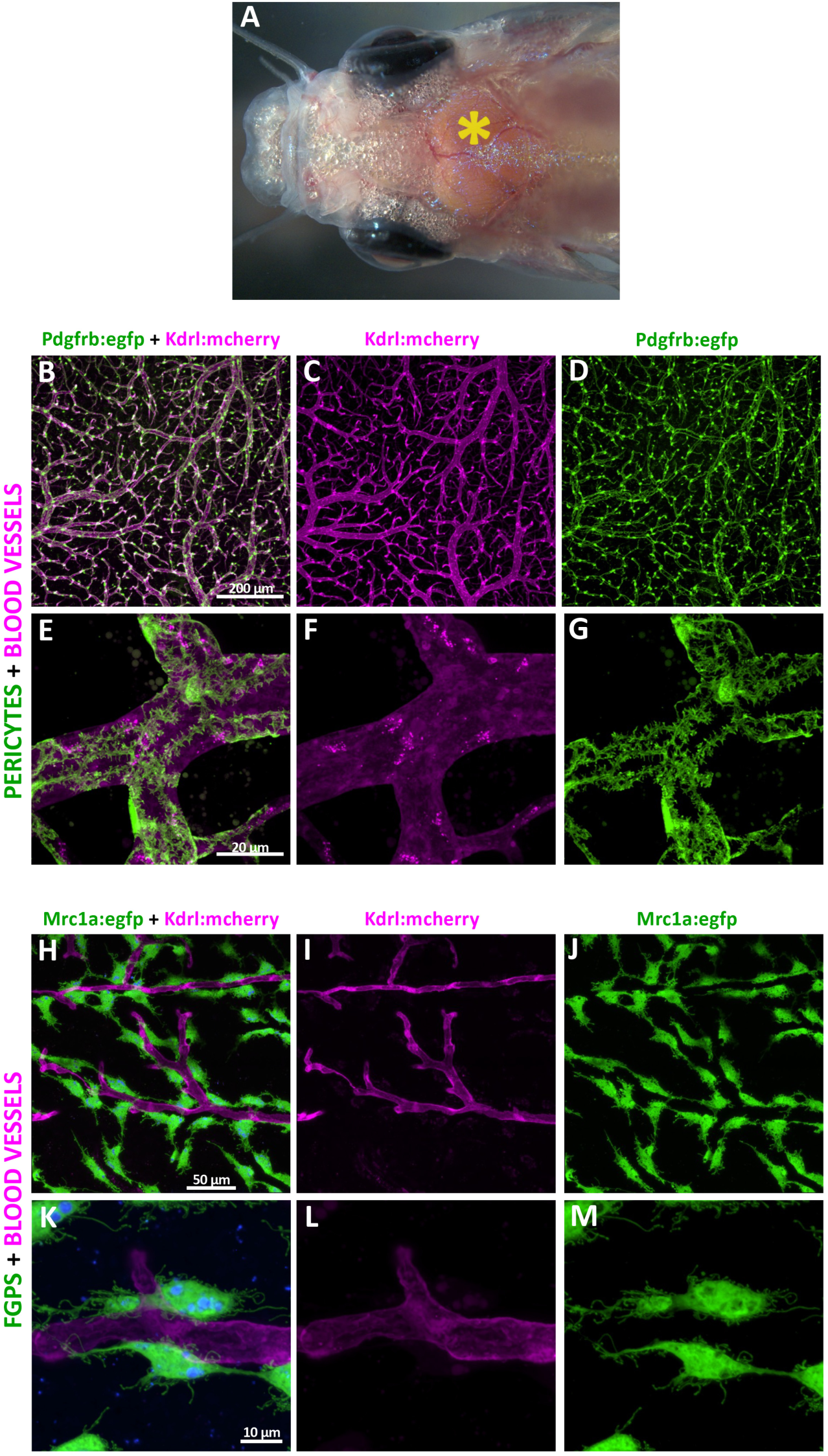
Vascular and vascular-associated cells in the leptomeninges. **A,** Dorsal view of a *casper* adult zebrafish head. The asterisk notes the optic tectum lobe area where the images in B-C were collected. **B-G,** Confocal images of EGFP-positive pericytes (B,D,E,G; green) and mCherry-positive blood vessel endothelial cells (B,C,E,F; magenta) in the leptomeninges of a dissected brain from a *Tg(kdrl:mcherry)^y206^*, *Tg(pdgfrb:egfp)^ncv22^*double transgenic adult zebrafish. Panels E-G show higher magnification images of blood vessels with pericytes. **H-M,** Confocal images of EGFP-positive FGPs (H,J,K,M; green) and mCherry-positive blood vessel endothelial cells (H,I,K,L; magenta) in the leptomeninges of a dissected brain from a *Tg(kdrl:mcherry)^y206^, Tg(mrc1a:egfp)^y251^ double transgenic adult zebrafish* double transgenic adult zebrafish. Panels K-M show higher magnification images of blood vessels with FGPs. Scale bars = 200 µm (B-D), 20 µm (E-G), 50 µm (H-J), and 10 µm (K-M).

**Supplemental Figure 5.**
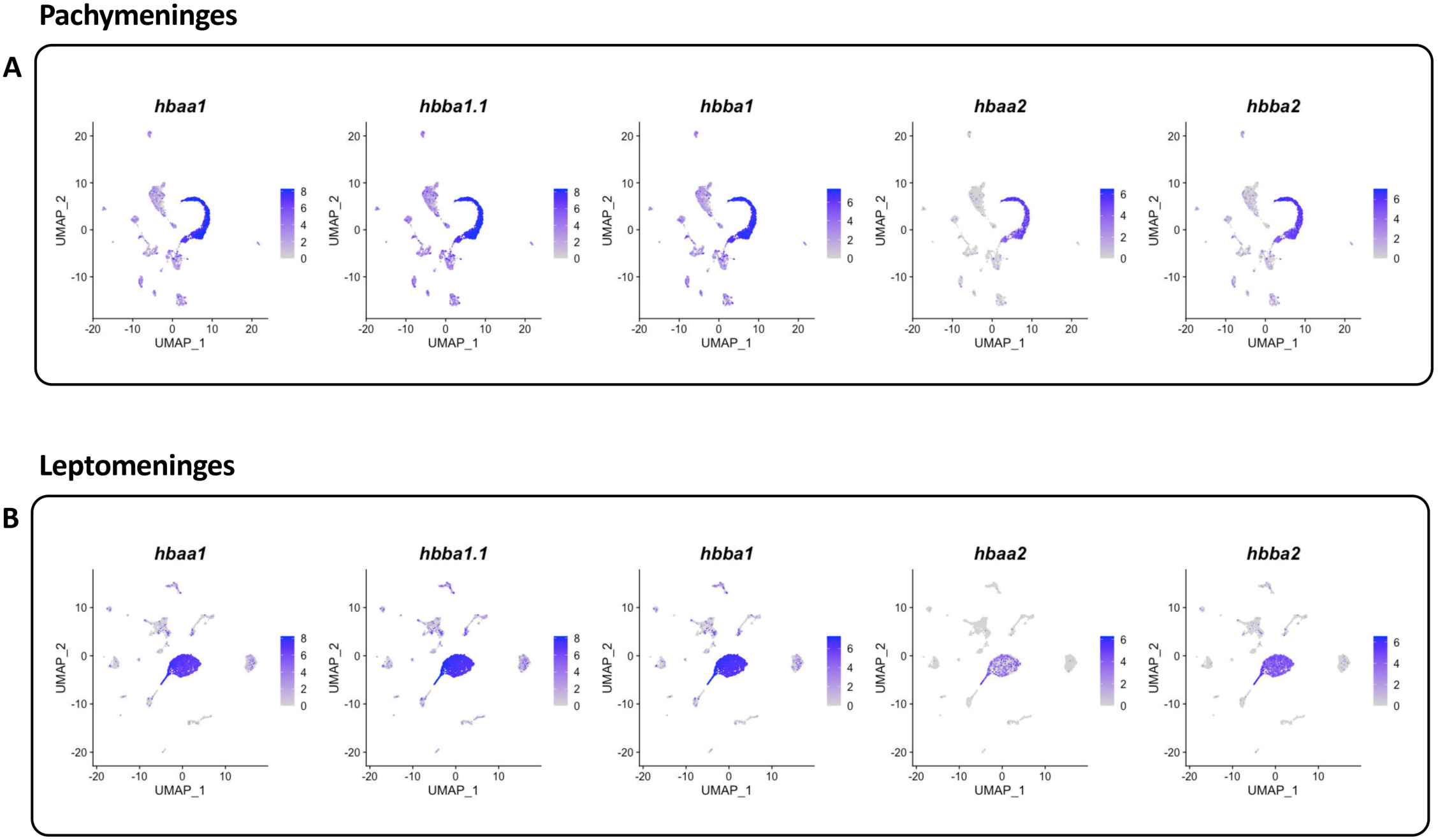
Identification of erythrocytes in meningeal scRNA-seq data sets. **A,** Pachymeningeal feature plots showing adult hemoglobins expressed in the pachymeningeal scRNA-seq data set, with very strong expression in the erythrocyte cluster. **B,** Leptomeningeal feature plots showing adult hemoglobins expressed in the leptomeningeal scRNA-seq data set, with very strong expression in the erythrocyte cluster.

**Supplemental Figure 6.**
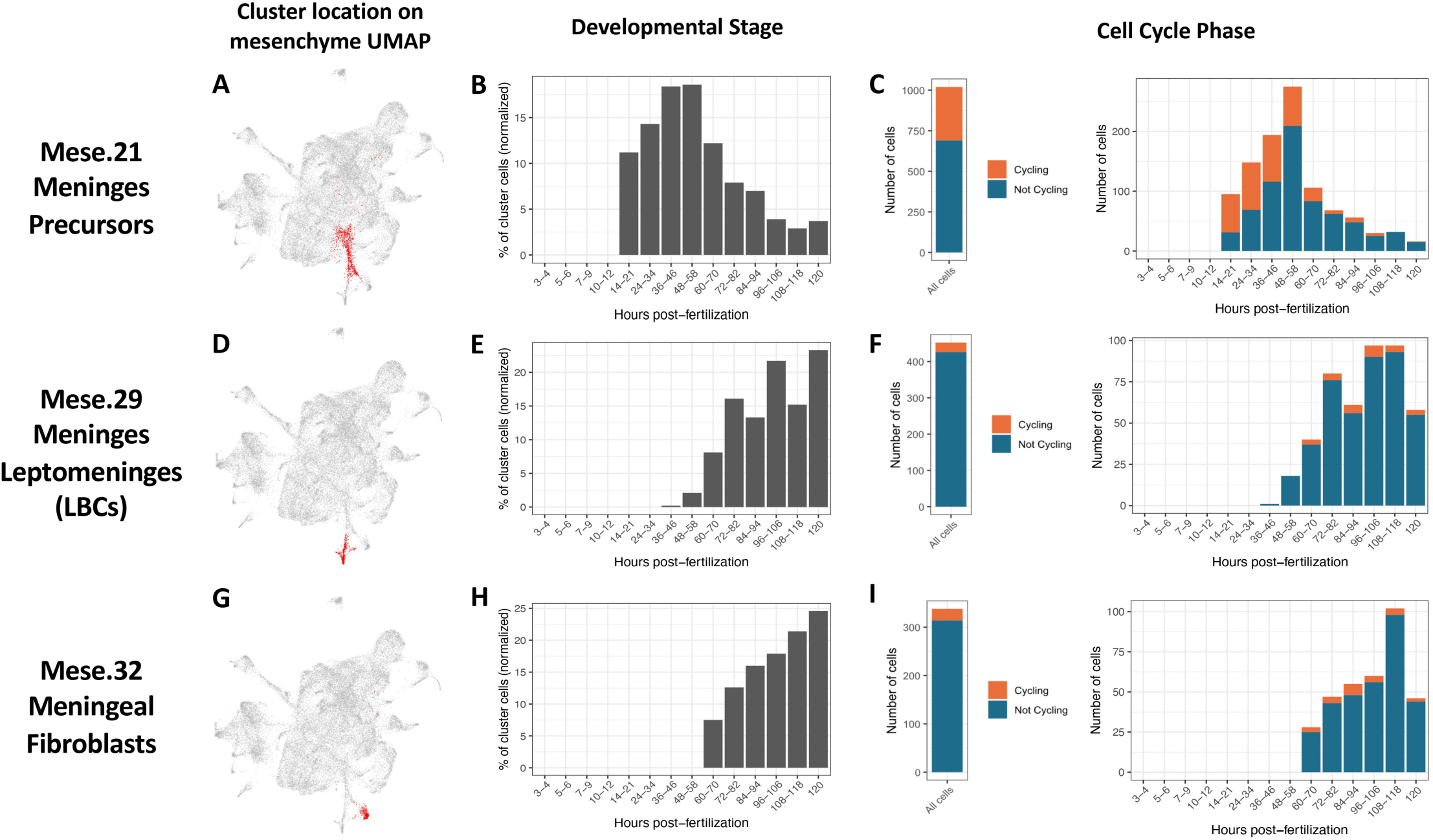
Meningeal precursor, meningeal fibroblast, and LBC clusters in the Daniocell mesenchyme scRNA-seq data set. **A,D,G,** UMAP plots of the mesenchymal subset of the Daniocell (https://daniocell.nichd.nih.gov/) scRNA-seq data set, highlighting the cells in each of the three clusters. **B,E,H,** Graphical plot of the normalized percentage of cells present at each developmental stage of the Daniocell data set, for each of the three clusters. **C,F,I,** Graphical plot of the number of cycling vs. non-cycling cells present at each developmental stage of the Daniocell data set, for each of the three clusters. Clusters shown in the panels are Mese.21 “Meninges Precursors” (**A-C**), Mese.29 “Meninges Leptomeninges” (**D-F**; e.g., LBCs), or Mese.32 “Meningeal Fibroblasts” (**G-I**). Mese.21 Meninges Precursors are situated at the base of both Mese.29 Meninges Leptomeninges and Mese.32 Meningeal Fibroblast cell clusters (A,D,G), and they appear at earlier stages than LBCs and meningeal fibroblasts (B,E,H), and include many more cycling cells, especially at the earlier developmental stages (C,F,I).

**Supplemental Figure 7.**
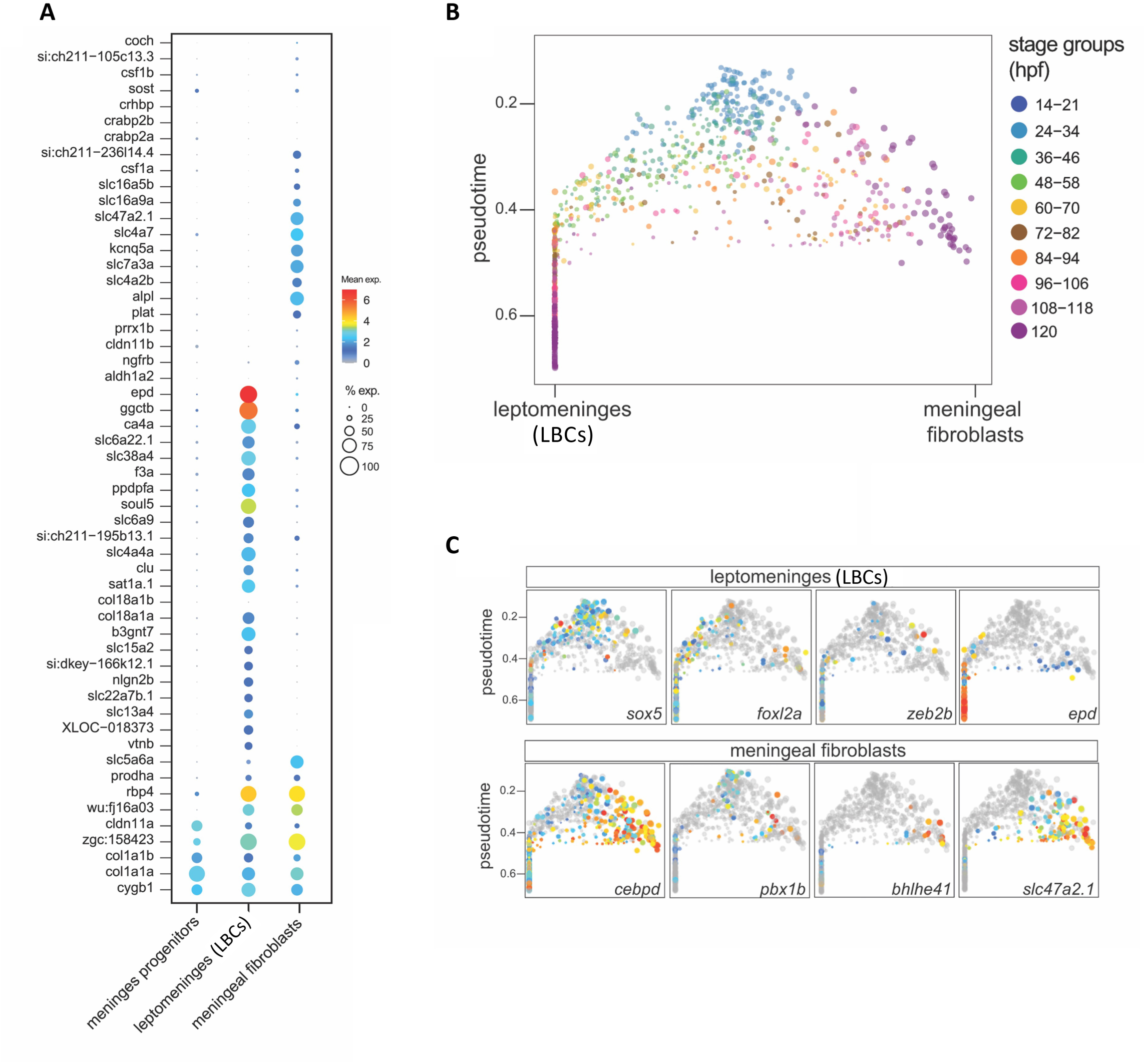
Supplementary data for LBCs vs. meningeal fibroblasts. **A,** Dot plot showing enriched expression of selected diagnostic genes in meningeal fibroblasts, LBCs, or both in the mesenchymal subset of the Daniocell scRNA-seq data set. Average expression and percent of cells in each cluster expressing each gene are indicated by dot color and size, respectively, with average expression on a log2 scale. **B,** Pseudotime plot of cells in the Mese.21 “Meninges Precursor,” Mese.29 “Meninges Leptomeninges,” and Mese.32 “Meningeal Fibroblast” clusters, with color coding showing the actual developmental stages of each cell. **C,** Pseudotime plots of cells in the Mese.21 “Meninges Precursor,” Mese.29 “Meninges Leptomeninges,” and Mese.32 “Meningeal Fibroblast” clusters, showing expression of genes enriched in either LBCs (top row) or meningeal fibroblasts (bottom row).

**Supplemental Figure 8.**
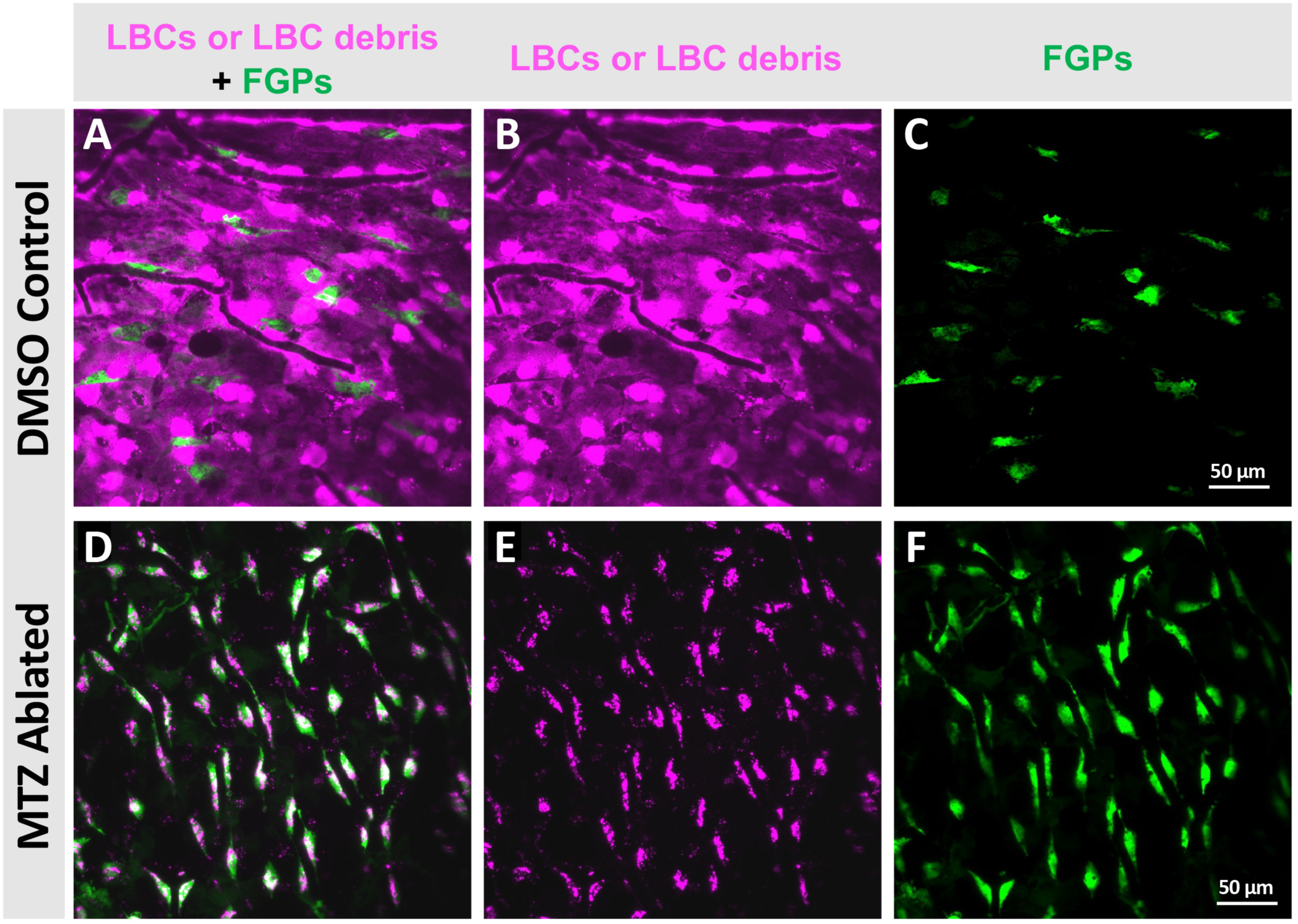
Apoptotic LBC cells are phagocytosed by adjacent FGPs after LBC ablation in the adult leptomeninges. **A-F.** Confocal micrographs of transgenic *Tg(epd:gal4-VP16);Tg(uas:ntr-mcherry);Tg(mrc1a:egfp)* adult animals treated with either DMSO (A-C) or 10mM MTZ (D-F). Top panels show a control DMSO treated animal with intact LBC (magenta) and FGP (green) leptomeningeal coverage. Bottom panels (D-F) show an animal with ablated LBCs, where FGPs (green) have engulfed the LBC-resulting debris (magenta) showing strong magenta inclusions inside them.

## SUPPLEMENTAL MOVIE LEGENDS

**Supplemental Movie 1. T cells, lymphatics, and FGPs in the meninges.** 3-D reconstructed views of a confocal Z-stack imaged through the skull of an intact, living *casper, Tg(lck:mcherry)^ns107^, Tg(mrc1a:eGFP)^y251^* double transgenic adult zebrafish, showing T cells (magenta) in and around lymphatic vessels (green) in the dural meninges to FGPs in the leptomeninges. Blue autofluorescence is used to visualize the skull located immediately above the meningeal layers.

**Supplemental Movie 2. LBCs on the surface of the larval zebrafish brain.** The first half of the movie shows a 14 hour time-lapse confocal imaging of LBC nuclei (magenta) and cell bodies (green) over the surface of an approximately 2-2.5 dpf *Tg(epd:mcherry)^y715^*, *Tg(epd:gfp-caax)^y716^* larval brain (selected frames shown in **Fig. 6H-L**). Yellow arrows show dividing LBCs. The second half of the movie shows rotating 3D reconstruction views of a confocal stack of LBCs on a 5 dpf *Tg(epd:gfp-caax)^y716^*brain, revealing that LBCs are located only on the surface of the brain (sequence begins with anterior to the left, dorsal in foreground).

**Supplemental Movie 3. The adult zebrafish leptomeninges is composed largely of LBCs.** 3-D reconstructed views of a confocal Z-stack through the leptomeninges on the surface of a dissected *Tg(epd:mCherry), Tg(mrc1a:eGFP)* double transgenic adult zebrafish brain, with LBCs in magenta and FGPs in green. Blood vessels are unlabeled but appear as black tubular structures surrounded by LBCs and FGPs. Movie begins with a dorsal (X-Y) view then rotates to show lateral (X-Z) slices with blood vessels and FGPs running through a continuous LBC layer. Scale bar = 25mm. Selected frames shown in **Fig. 6P,Q**).

**Supplemental Movie 4. Metronidazole-dependent LBC ablation in larval zebrafish.** Video showing lateral views of 5 dpf transgenic *Tg(epd:gal4-VP16);Tg(uas:ntr-mcherry);Tg(mrc1a:egfp)* zebrafish larvae treated with either 10mM MTZ (upper panel) or DMSO (lower panel) showing MTZ-NTR dependent induction of LBC apoptosis. Time lapse was started at 5 dpf with acquisition intervals every 10 minutes for a total of 12 hours.

**Supplemental Movie 5. Control and LBC-ablated adult zebrafish.** Video images of the gross behavioral effects of acute LBC ablation on adult zebrafish. Control DMSO carrier only treated (LBCs intact) animals are in the lower tank, and 10mM MTZ treated (LBCs ablated) animals are in the upper tank. Animals are shown 12 hours post-initiation of treatment (comparable to the animals shown in **Fig. 8O-V**). Control animals are unaffected, but ablated animals are unresponsive and swimming slowly or unable to swim. Most EDP-ablated animals are still alive (possessing an active heartbeat) at this stage, but all will die within hours.

**Supplemental Movie 6. Acute ablation of LBC cells in adult animals.** Video shows 3D renderings of confocal micrographs of either control (left) or MTZ-treated adult *Tg(epd:gal4-VP16);Tg(uas:ntr-mcherry);Tg(mrc1a:egfp)* transgenic animals following exposure to DMSO or 10mM MTZ for 24 hours. Images were obtained on alive animals through intact skulls to preserve meningeal anatomy. LBCs are shown in magenta, FGPs and lymphatics are in green and skull (autofluorescence) is shown in blue. MTZ-treated animal shows apoptotic LBCs (magenta) being phagocytosed by FGPs (green).

## SUPPLEMENTAL FILES

Supp Figs S1-S8: Venero Galanternik et al – Supp Figures 1-8.pdf

Supplemental Movie 1. Venero_et_al_Supp_Movie_1.mp4

Supplemental Movie 2. Venero_et_al_Supp_Movie_2.mp4

Supplemental Movie 3. Venero_et_al_Supp_Movie_3.mp4

Supplemental Movie 4. Venero_et_al_Supp_Movie_4.mp4

Supplemental Movie 5. Venero_et_al_Supp_Movie_5.mp4

Supplemental Movie 6. Venero_et_al_Supp_Movie_6.mp4

---

**SUPPLEMENTAL TABLE 2.**
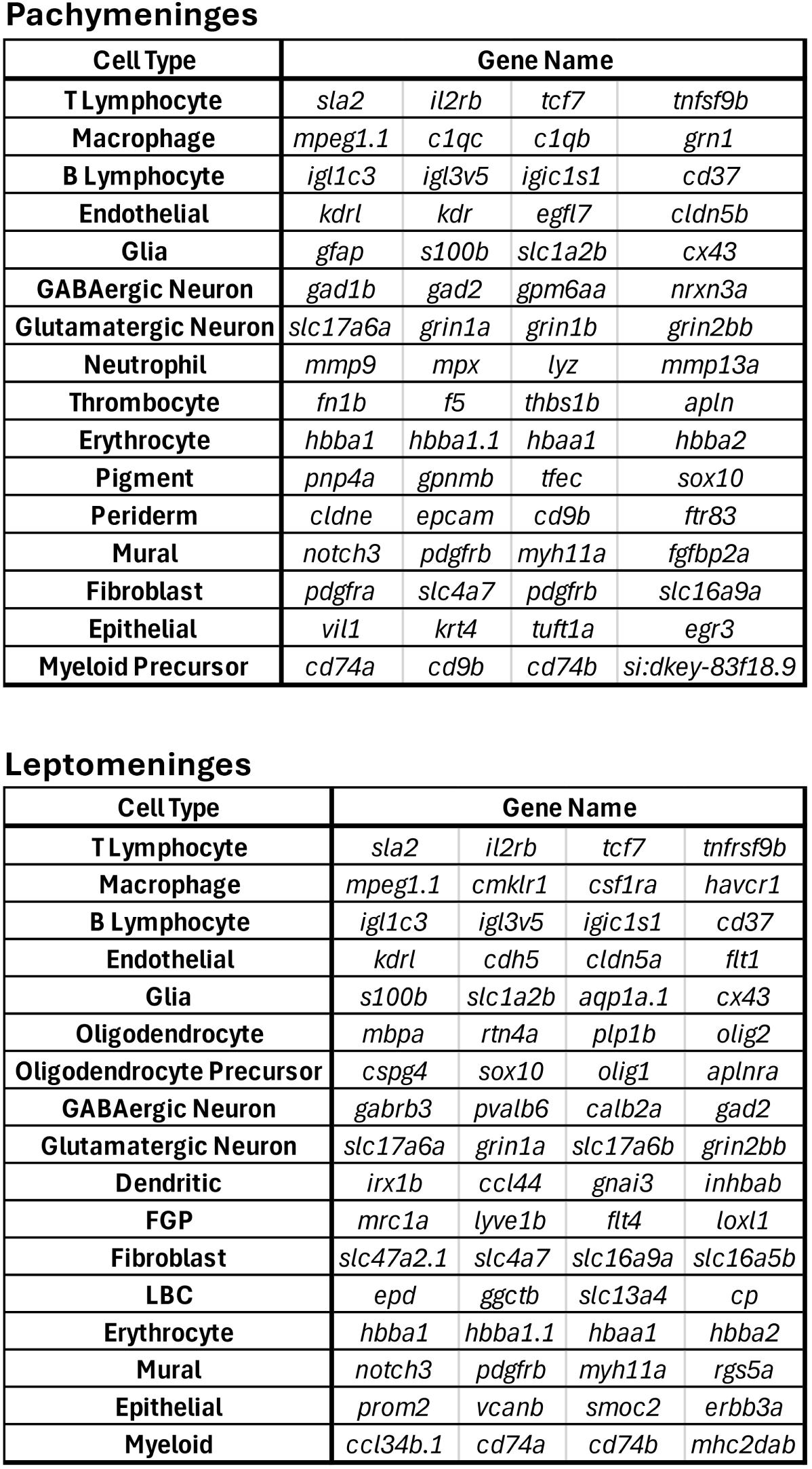

